# Alterations in the gut microbiome by HIV-1 infection or a high-fat diet associates with systemic immune activation and inflammation in double humanized-BLT mice

**DOI:** 10.1101/2020.03.16.993956

**Authors:** Lance Daharsh, Amanda E. Ramer-Tait, Qingsheng Li

## Abstract

**Background:** While the translatability of gut microbiome studies utilizing animal models to humans has proven difficult, studying the gut microbiome directly in humans is also challenging due to the existence of many confounding variables. Therefore, we utilized double humanized mice, which have both an engrafted stable human-like gut microbiome and functional human immune system. With this model, we were able to determine the in vivo impact of HIV-1 infection or a high-fat diet (HFD) on gut human microbiome composition, and its relationship with human immune cell activation and systemic inflammation.

**Results:** Surgery was performed on NSG mice to create humanized bone-marrow, liver, thymus mice (hu-mice). In order to create double hu-mice, the hu-mice were treated with broad spectrum antibiotics to deplete murine gut bacteria and subsequently transplanted with human fecal material from healthy human donors. We characterized 262 fecal samples from hu-mice, double hu-mice, and human fecal donors to determine the impact of HIV-1 infection or HFD on the gut microbiome and systemic immune activation and inflammation. We found that HIV-1 infection altered the human-like gut microbiome of double hu-mice, which was associated with decreased human CD4 T cells and increased systemic inflammation and immune activation. Further, using a HFD we induced gut microbial dysbiosis in double hu-mice which corresponded with increased systemic immune activation and inflammation.

**Conclusions:** Here, we describe the changes in the human gut microbiome and human immune system due to HIV-1 infection or HFD using our double hu-mice model. HIV-1 infection led to changes in the composition of the human-like gut microbiome that was associated with human CD4 T cell loss and high levels of inflammation and immune activation. The HFD quickly changed the composition of the gut microbiome and led to systemic immune activation and inflammation. We further identified a subset of gut bacteria in HIV-1 infected and HFD fed double hu-mice that was closely associated with systemic inflammation and immune activation. This study demonstrated how double humanized mice can be used to study the complex in vivo interactions of the gut microbiome and human immune system in the context of both disease and diet.

## Background

The human gut is home to the largest number of immune cells in the body and provides an ecosystem for trillions of microbes known collectively as the gut microbiome [1, 2]. The gut microbiome and corresponding gut immune system have a highly reciprocal and dynamic relationship that is a critical determinant for human health and disease [3-7]. While the gut microbiome influences host immune responses through their antigens and metabolites, the immune system in turn contributes to shaping the composition and distribution of gut microbes [8-10]. It has been estimated that up to 10% of immune response variability is associated with the gut microbiome [11]. The gut microbiome has also been shown to be essential for proper immune development, immune function, and response to infection and vaccination.

The gut plays a key role in the pathogenesis of human immunodeficiency virus type-1 (HIV-1) infection. One of major features of HIV-1 infection is the rapid and extensive loss of gut immune cells, most notably CD4+ T cells [12-17]. The depletion of gut immune cells is accompanied by alterations in the gut microbiota, the interruption of gut epithelial barrier integrity with subsequent microbial translocation, and increased inflammation and immune activation [18-25]. In addition, the alterations of the gut microbiome and gut immune cells may increase the susceptibility to rectal HIV-1 transmission, as previous studies have shown that local and systemic inflammation will activate and attract CD4+ T cells thereby increasing the risk of HIV-1 mucosal transmission [26]. Further, gut microbial dysbiosis and microbial translocation have been implicated in the incomplete gut immune reconstitution and increased systemic immune activation and inflammation in people living with HIV (PLH) on suppressive anti-retroviral therapy (ART) [21, 25, 27-29]. Despite the ability of ART to suppress HIV-1 replication to undetectable levels in peripheral blood, gut immune reconstitution following ART is often slow and incomplete [30-35]. Additionally, there are persistently increased levels of inflammation and immune activation [36-38] that contribute to increased comorbidities in PLH [39-41], of which gut microbial translocation mediated immune activation is thought to be a major contributing factor [42, 43]. Consequently, in the ART era, the life expectancy of HIV-1 infected individuals in developed countries is over 10-years shorter than a normal lifespan [44] and non-infectious morbidities are also significantly higher than the general population [45]. Given the importance of the gut to almost all aspects of prevention, pathogenesis, and treatment of HIV-1, investigating the in vivo relationship between the gut microbiome and human immune system may provide novel insights for prevention and treatment strategies. Due to the bidirectional relationship of the gut microbiome and the immune system, resolving gut microbial dysbiosis may also improve immune cell recovery, reduce immune activation and inflammation during ART treatment, and ultimately reduce comorbidities.

Despite the significant progress that has been made in understanding HIV-1 pathogenesis and in treating HIV-1 infection, a key knowledge gap remains in our mechanistic understanding of the impact of the gut microbiome on immune activation and inflammation during HIV-1 infection. Research utilizing animal models has provided a large portion of our understanding of the connection between the gut microbiome and human disease, of which humanized mice (hu-mice), that feature an engrafted human immune system, are an important pre-clinical animal model for translational biomedical research [46-54]. However, the gut murine microbiome significantly differs from humans due to anatomical, evolutional, environmental, and diet differences [55, 56]. To improve the translatability of hu-mice research, we previously developed double hu-mice that feature a stable human-like gut microbiome and human immune system [57, 58]. In this study, we used this model and investigated the immunopathogenesis of alterations in the gut microbiome induced by HIV-1 infection and a high-fat diet (HFD). We found that HIV-1 infection led to changes in the composition of the human-like gut microbiome that temporally corresponded with human CD4+ T cell loss and high levels of inflammation and immune cell activation. We also showed that a HFD quickly changed the composition of the gut microbiome and led to systemic immune activation and inflammation. Importantly, this study demonstrated the double hu-mice model can be used to study the complex in vivo interactions of the gut microbiome and human immune system in the context of human health and disease.

## Results

### HIV-1 infection altered the gut microbiome of hu-BLT mice

The hu-BLT mice (hu-mice) model allows for a high level of human immune cell reconstitution and has been used for the study of HIV-1 prevention, pathogenesis, and treatment [59-62]. However, the hu-mice gut microbiome is of murine origin and we previously demonstrated that hu-mice harbored a distinct low diversity gut microbiome [63]. To investigate the extent to which HIV-1 infection impacts the murine gut microbiome in this model, we compared longitudinally sampled gut microbiome profiles of HIV-1 infected hu-mice and uninfected hu-mice. Twenty-four fecal samples from 4 HIV-1 infected hu-mice were collected longitudinally for up to 6 consecutive weeks. The HIV-1 infected hu-mice had an altered gut microbiome composition compared to uninfected hu-mice and the two groups clustered distinctly from one another in both Non-metric Multi-dimensional Scaling (NMDS) and Principal Coordinates Analysis (PCoA) plots (Supplemental Figure HIV_HuMice.pdf). Additionally, HIV-1 infected hu-mice had higher measures of alpha diversity, including the number of unique species per sample or species richness, Simpson’s Diversity Index, and Shannon Diversity Index (Supplemental Figure HIV_HuMice.pdf). There were multiple alterations in the gut microbiome composition of the HIV-1 infected hu-mice as shown by heatmaps of bacterial relative abundance for Order and Family taxa levels (Supplemental Figure HIV_HuMice.pdf). Significant differences in the relative abundance of gut bacterial taxa were found using Kruskal-Wallis tests with false discovery rate (FDR) adjusted P values <.05 (Supplemental File HIV_HuMice_KW.xlxs). Nevertheless, due to the large compositional differences between the murine and human gut microbiomes, regular hu-mice without a humanized gut microbiome may limit their translatability for the study of human health and disease [55, 56].

### HIV-1 infection altered the gut microbiomes of double humanized BLT mice

We previously developed a double hu-mice model that harbors a stable human-like gut microbiome in addition to a functional human immune system [58, 63]. To create double hu-mice, we first performed surgery on NSG mice to create hu-BLT mice with an engrafted human immune system. Hu-mice were subsequently treated with a cocktail of broad-spectrum antibiotics to reduce murine gut bacteria, followed by human fecal material transplants (FMT) using fecal material from a mixture of three healthy human donors. This double hu-mice model was used to determine if and how the composition of the gut microbiome was altered during HIV-1 infection (Table 1). Gut microbiome profiles of the mice were sampled at 10 weeks post BLT humanization surgery and before antibiotic treatment and FMTs (Pre-FMT) as well as one week following the completion of antibiotic treatment and FMTs (Post-FMT).

**Table 1.** Summary of experimental hu-mice cohorts.

| Cohort | Double hu-mice | Number of mice | Number of fecal samples | Condition | Maximum weeks collected |
| --- | --- | --- | --- | --- | --- |
| Pre-FMT | No | 40 | 40 | NA | 1 |
| Post-FMT | Yes | 32 | 31 | NA | 1 |
| HIV infected double hu-mice | Yes | 14 | 90 | HIV-1 | 12 |
| HFD | Yes | 8 | 34 | High fat diet | 7 |
| LFD | Yes | 8 | 39 | Low fat diet | 9 |

After collecting the Post-FMT fecal samples, double hu-mice were intraperitoneally injected with 4.5*10^5 TCID of an equal mixture of HIV-1_SUMA_ and HIV-1_JRCSF_. To determine the longitudinal changes to the gut microbiomes of HIV-1 infected double hu-mice, fecal samples were collected every week for up to 12 weeks from 11 infected double hu-mice (Infected) and 3 uninfected double hu-mice (Uninfected). As shown in Figure 1AB, HIV-1 infection altered the composition of the gut microbiome over the course of the study. Additionally, sample collection date (Figure 1CD) and body weight (Figure 1 EF) were associated with the composition changes observed in the gut microbiome.

**Figure 1.**
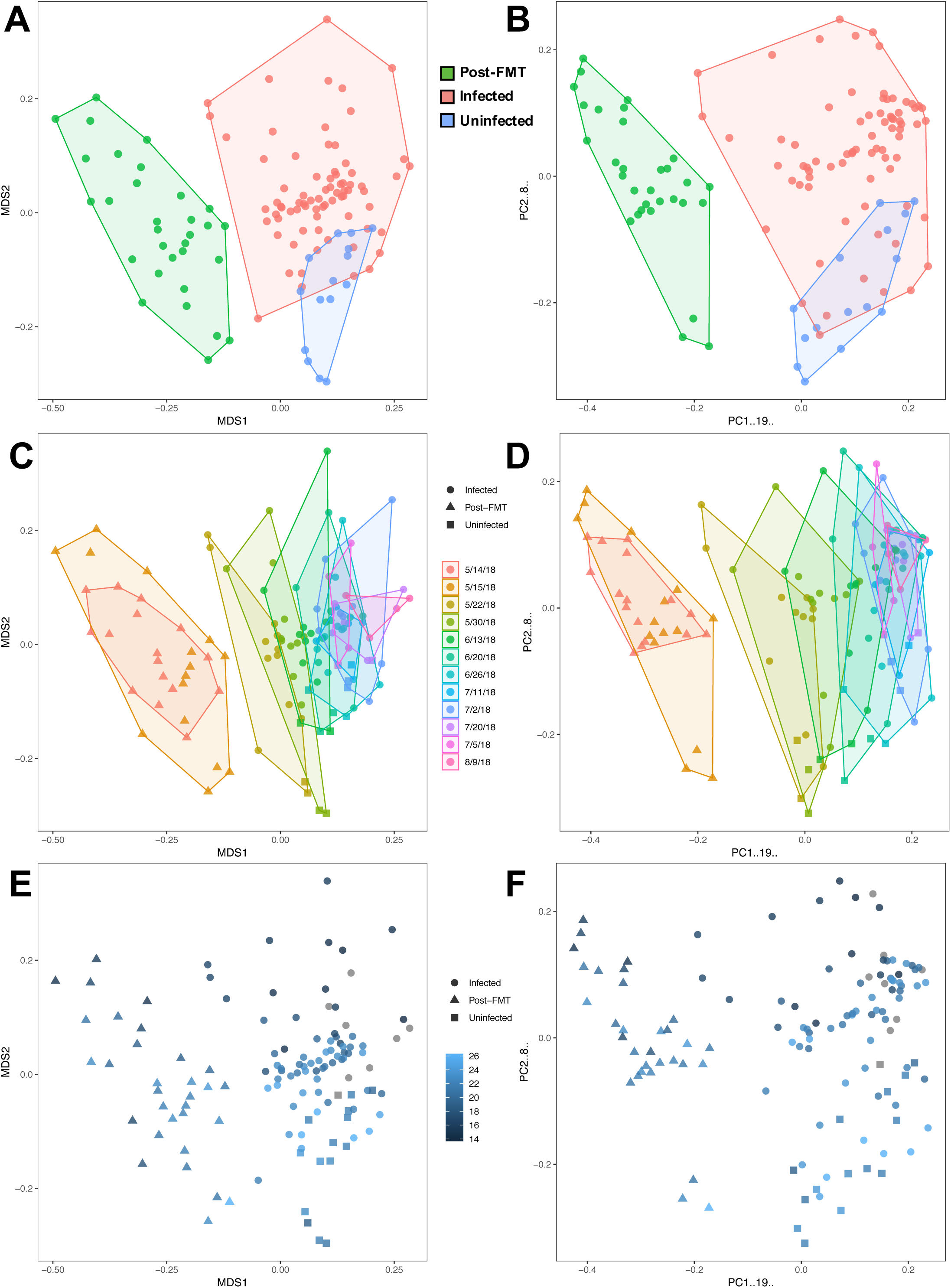
HIV-1 infected double hu-mice had a significantly different gut microbiome composition compared to uninfected double hu-mice. Non-metric multidimensional scaling (NMDS) and Principal coordinate analysis (PCoA) plots AB) Clustering of post fecal material transplant (Post-FMT), HIV-1 infected (Infected) and uninfected (Uninfected) double humanized mice gut microbiome profiles. CD) Clustering of post fecal material transplant (Post-FMT), HIV-1 infected (Infected) and uninfected (Uninfected) double humanized mice gut microbiome profiles based on sample collection date. EF) Clustering of post fecal material transplant (Post-FMT), HIV-1 infected (Infected) and uninfected (Uninfected) double humanized mice gut microbiome profiles based on body weight on collection date.

Double hu-mice form both the Infected and Uninfected group had altered gut microbiome profiles compared to Post-FMT samples (Figure 2AB). Both infected and uninfected double hu-mice had slightly higher measures of alpha diversity compared to Pre-FMT and Post-FMT samples, including the number of unique species per sample or species richness, Simpson’s Diversity Index, and Shannon Diversity Index (Figure 2CDE). Infected samples had slightly higher alpha diversity measures compared to uninfected samples, but the differences were not significant. Differences in relative abundance for the experimental groups are shown by Order and Family taxa levels (Figure 2FG). Significant differences in the relative abundance of gut bacterial taxa between the double hu-mice from the Infected group and Uninfected group were found using Kruskal-Wallis tests with false discovery rate (FDR) adjusted P values <.05 (Supplemental File HIV_KW.xlsx). Infected double hu-mice had a higher relative abundance of *Bifidobacteriaceae* (7.09%, P=0.0001 FDR) and *Ruminococcaceae* (4.47%, P=0.0005 FDR) and a lower abundance of *Lactobacillaceae* (−15.67%, P=0.0001 FDR) and *Turicibacteraceae* (−3.99%, P=0.0302 FDR) (Supplemental File HIV_Composition.pdf).

**Figure 2.**
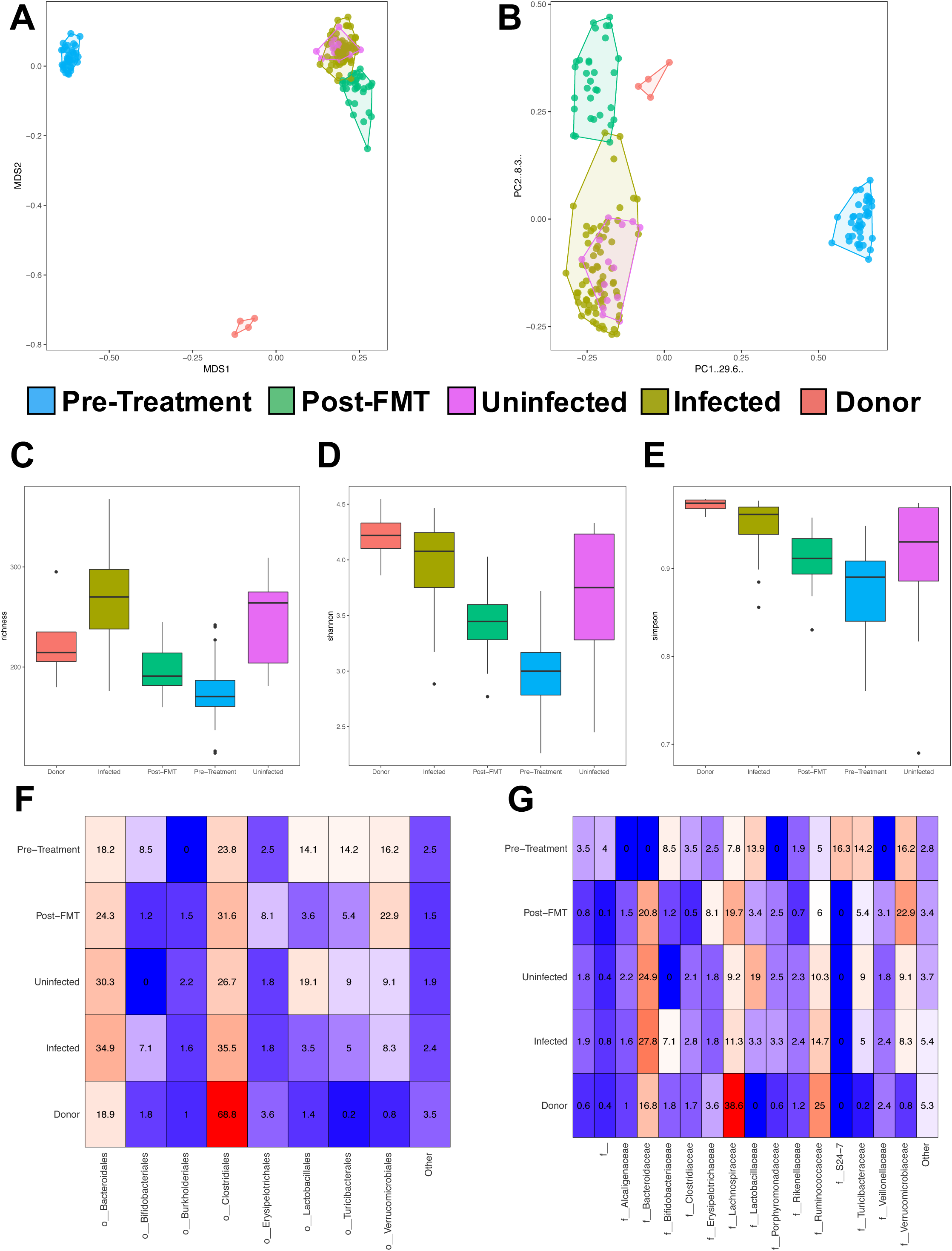
HIV-1 infected double hu-mice had a significantly different gut microbiome composition compared to uninfected double hu-mice. Gut microbiome profiles for humanized mice before receiving antibiotic treatment and subsequent fecal material transplants (Pre-treatment), double humanized mice post fecal material transplant (Post-FMT), HIV-1 infected double humanized mice (Infected) and uninfected double humanized mice (Uninfected), and human donor fecal samples (Donor) A) Gut microbiome profiles displayed by Non-metric multidimensional scaling (NMDS). B) Gut microbiome profiles displayed by Principal coordinate analysis (PCoA). C) Alpha diversity of gut microbiome profiles shown by species richness. D) Alpha diversity of gut microbiome profiles shown by Shannon Index. E) Alpha diversity of gut microbiome profiles shown by Simpson Index. F) Taxa abundance plot of gut microbiome profiles by Order level. G) Taxa abundance plot of gut microbiome profiles by Family level.

To determine the changes in the gut microbiome of double hu-mice before and after HIV-1 infection, we compared infected samples with Post-FMT samples (Supplemental File HIV_KW.xlxs). After infection, double hu-mice had a lower relative abundance of *Erysipelotrichaceae* (−6.18%, P=0.0001 FDR), *Lachnospiraceae* (−4.96%, P=0.0051 FDR), and *Verricomicrobiaceae* (−11.82%, P=0.0001 FDR) and a higher relative abundance of *Bacteroidaceae* (4.52%, P=0.0424 FDR), *Bifidobacteriaceae* (4.43%, P=0.0404 FDR), *Clostridiaceae* (2.48%, P=0.0001 FDR), *Rikenellaceae* (1.63%, P=0.0001 FDR), and *Ruminococcaceae* (8.64%, P=0.0001 FDR).

A random forest model was trained to predict if the gut microbiome profiles came from double hu-mice that were HIV-1 infected or uninfected based on the amplicon sequence variant (ASV) features. The top 15 most important discriminatory features of the model based on area under the ROC curve were then identified. These features were scaled to 100 and plotted along with the average normalized ASV counts for each group (Supplemental File HIV_Importance.pdf). The top ranked features were ASV527 *Ruminococcus* and ASV321 *Dorea*, both of which were more prevalent in infected double hu-mice. Of the top 15 features, many ASVs were more prevalent in HIV-1 infected mice including ASVs from *Butyricicoccus pullicaecorum, Ruminococcaceae, Ruminococcus, Oscillospira*, and *Christensenellaceae*. Three ASVs were more prevalent in uninfected double hu-mice including ASVs from *Butyricicoccus pullicaecorum, Blautia producta*, and *Lachnospiraceae*. Here we show there are many features of the gut microbiome that were different between infected and uninfected double hu-mice. To further evaluate the role of gut microbiome during HIV-1 infection we determined the inflammatory and immune profiles from these double hu-mice.

### HIV-1 infection of double hu-mice led to increased systemic inflammation and immune activation

To evaluate systemic inflammation and immune activation in HIV-1 infected and uninfected double hu-mice, we measured plasma proinflammatory cytokines using multiplex immunoassays and human T cell activation in peripheral blood and splenic tissues using flow cytometry. Infected double hu-mice had significantly higher levels of IL-1β, IL-6, IFN-γ, and TNF-α (Figure 3) at 12 weeks post infection (WPI), whereas, cytokine levels from samples collected at 7 (WPI) were not elevated compared to uninfected samples. CD4 T cell depletion is the pathogenic hallmark of HIV-1 infection. HIV-1 infection leading to CD4 T cell death can be observed in peripheral blood of infected hu-mice beginning at 2-3 WPI [64]. Using flow cytometry, we tracked peripheral blood CD4 T cell levels in infected and uninfected double hu-mice (Figure 4AB). There was a decline in CD4 T cells in all post infection samples, of which some infected double hu-mice declined to levels below 50% of parent gated CD3 T cells. We also measured markers of immune activation in peripheral blood human T cells. All three populations of activated CD8 T cells, including CD8+ CD38+, CD8+ CD69+, and CD8+ HLA-DR+ T cells, were increased as a result of infection. CD4+ HLA-DR+ populations were also increased after infection, while the CD4+ CD69+ population had no significant changes. CD4+ CD38+ populations decreased after infection, which may be due to increased cell death in this population of activated CD4+ T cells.

**Figure 3.**
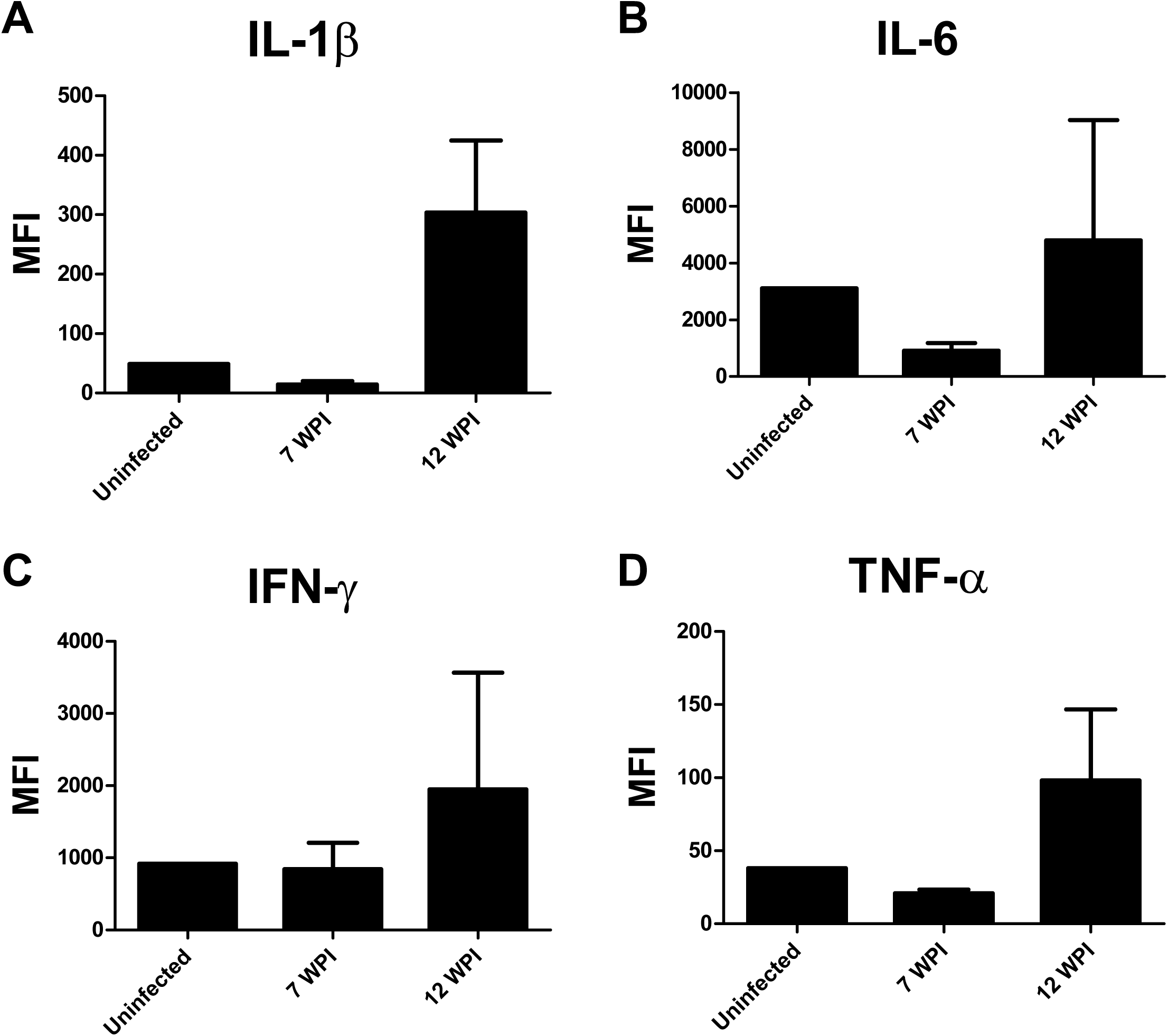
HIV-1 infected double hu-mice had increased systemic human inflammatory cytokines. Human inflammatory cytokine measures from plasma of double humanized mice. Samples were collected from double humanized mice at 7 and 12 weeks post infection (WPI). Cytokine levels shown by mean fluorescence intensity (MFI). A) IL-1β B) IL-6 C) IFN-γ D) TNF-α.

**Figure 4.**
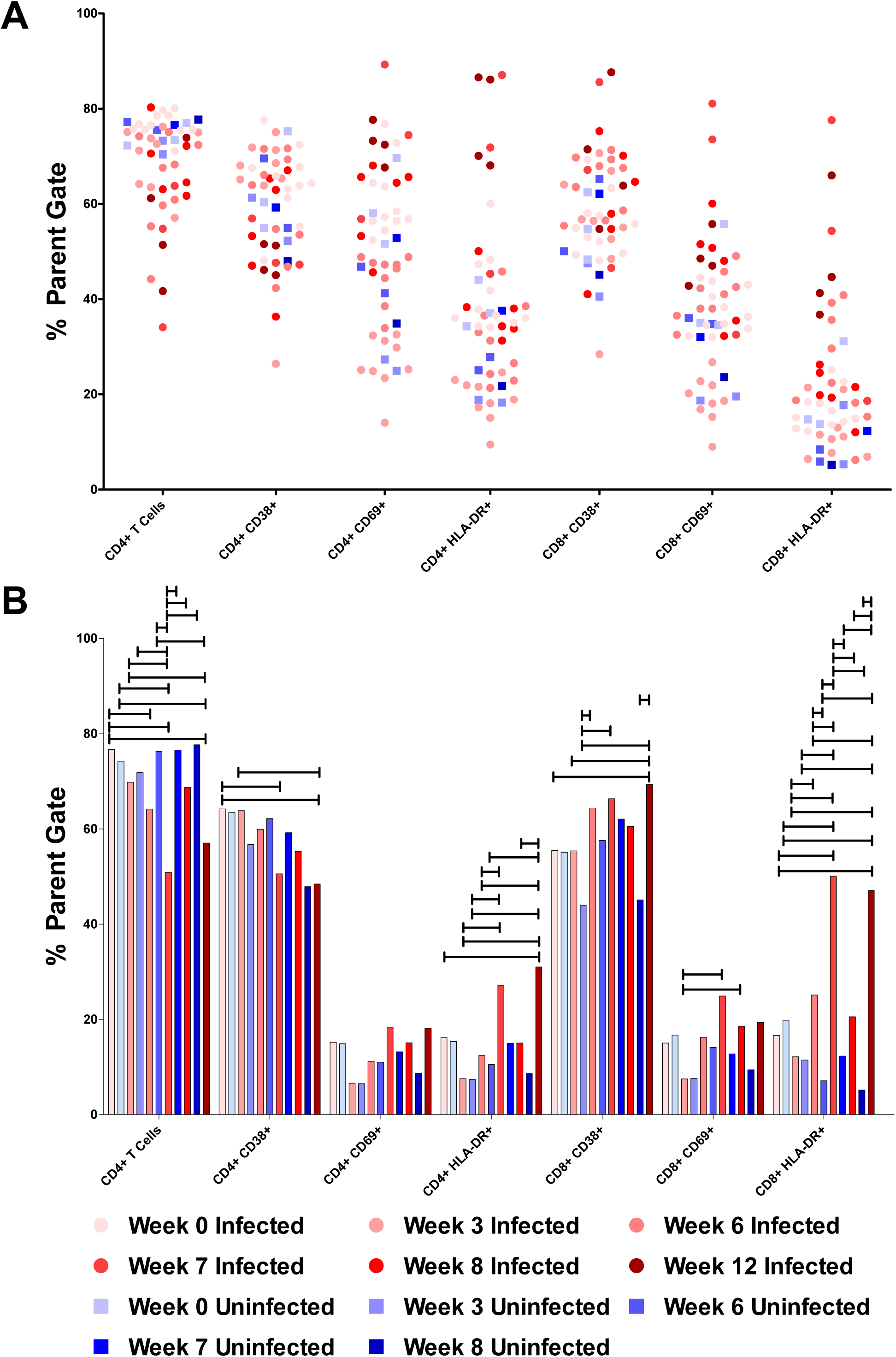
HIV-1 infected double hu-mice had increased systemic human immune cell activation. A) Human immune cell populations from peripheral blood of double humanized mice up to 12 weeks post HIV-1 infection. All immune populations were lymphocyte+, human CD45+ and mouse CD45-, and human CD3+) and are represented by the percentage of their parent gate. B) Percentage of peripheral blood human immune cell populations shown as a mean for the longitudinally collected HIV-1 infected or uninfected double humanized mice. For each population of human immune cells a multiple comparison test for significance was performed between the sample groups (ANOVA with Tukey Test with adjusted P values < 0.05).

During HIV-1 infection the level of CD4 T cell death and immune activation can differ between peripheral blood and lymphoid tissues. Therefore, 4 double hu-mice were sacrificed at both 7 WPI and 12 WPI. Flow cytometry was performed on lymphocytes isolated from spleen tissue (Supplemental Figure HIV_Spleen.pdf). The CD4 T cell loss was more severe in the spleen as compared to peripheral blood. There did not appear to be any major changes in CD4+ CD38+ population, while CD4+ CD69+ and CD4+ HLA-DR+ populations were increased in some of the infected animals. Almost all of the Infected samples had higher proportions of immune activated CD8 T cells, including CD8+ CD38+, CD8+ CD69+, and CD8+ HLA-DR+ populations. The double hu-mice model of HIV-1 infection recapitulates many important aspects of HIV-1 pathogenesis, including CD4 T cell loss and increased systemic inflammation and immune activation in the context of a human gut microbiome.

### Gut microbial dysbiosis was established in the double hu-mice model with a high-fat diet

The establishment of gut microbial dysbiosis in the double hu-mice model is needed for the investigation into the role of the gut microbiome in HIV-1 rectal transmission susceptibility and the study of the increased risk of comorbidities in HIV-1 infected individual on antiretroviral therapy. Previous studies have shown that local and systemic inflammation, along with the availability and activation state of target cells, are the major factors in determining the risk for HIV-1 transmission [26]. Studies on vaginal HIV-1 transmission demonstrated that the mucosal microbiome plays an important role in determining HIV-1 susceptibility [65-67]. Previous studies showed that feeding mice a high-fat diet (HFD) resulted in microbial dysbiosis, disruption of the gut epithelial barrier, increased systemic inflammation, and higher numbers of activated immune cells [68, 69]. Therefore, we fed double hu-mice with a HFD and found that it changed the engrafted healthy human gut microbiome into a state of microbial dysbiosis. The HFD group (N=8) was fed a diet consisting of 60% kcal from fat with 275 kcal of added sucrose (D12492, Research Diets Inc.). The low-fat diet (LFD) group (N=8) was fed a matched calorie control diet with 10% kcal from fat and no added sucrose (D12450K, Research Diets Inc.). Using this experimental design, we determined the impacts of these different diets on the gut microbiome as well as systemic inflammation and immune activation (Table 1). Before the introduction of a HFD or LFD, double hu-mice were fed regular mouse chow containing at least 14% protein (Teklad 2914). Fecal samples were collected for up to 9 weeks post new diet introduction.

The HFD group quickly showed drastic changes in gut microbiome composition as compared to Post-FMT samples of double hu-mice fed with regular mouse chow and double hu-mice fed with a LFD (Figure 5AB). The microbiome profiles from fecal samples collected from HFD and LFD fed groups clustered separately from the regular mouse chow Post-FMT samples based on principal component 1 (PC1). Further, the microbiome profiles from fecal samples collected from the HFD and LFD fed groups clustered separately from one another based on principal component 2 (PC2). After Pre-FMT and human donor samples were added to the analysis, the HFD and LFD samples clustered distinctly from the regular mouse chow Post-FMT samples (Supplemental Figure Diet_Comp.pdf). In the PCoA plot, PC1 represent the differences observed between the pre-existing murine gut microbiome in the Pre-FMT samples compared to the human-like gut microbiomes in double hu-mice and human donor samples. PC2 shows the differences between the regular mouse chow Post-FMT samples compared to the HFD and LFD samples.

**Figure 5.**
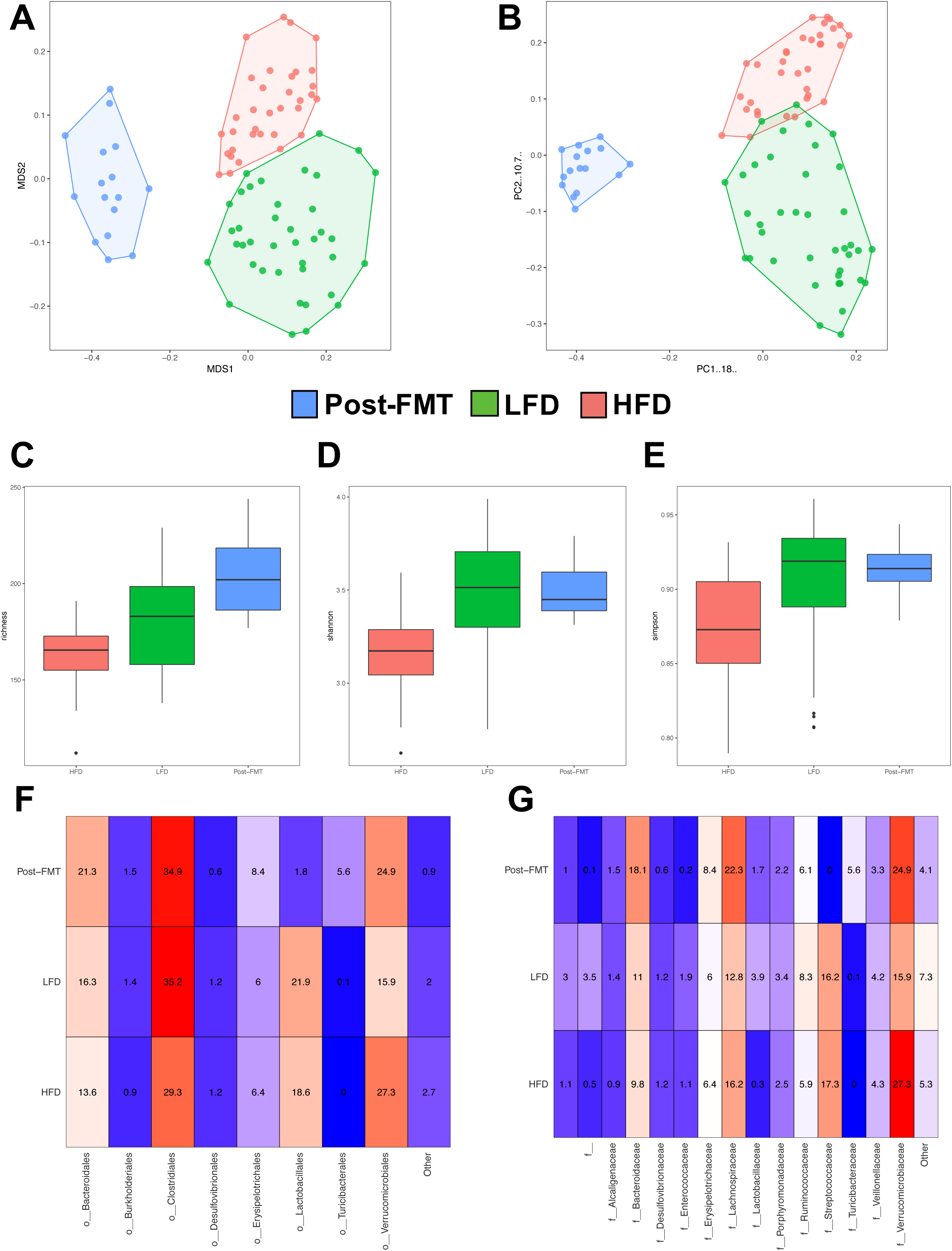
Diet significantly altered the gut microbiome of double hu-mice. A) Non-metric multidimensional scaling (NMDS) plot displaying double humanized mice on a mouse chow diet, a high fat diet, or a low fat diet. B) Principal coordinate analysis (PCoA) displaying double humanized mice on a mouse chow diet, a high fat diet, or a low fat diet. C) Alpha diversity plot of species richness comparing double humanized mice on a mouse chow diet, a high fat diet, or a low fat diet. D) Alpha diversity plot of the Shannon index comparing double humanized mice on a mouse chow diet, a high fat diet, or a low fat diet. E) Alpha diversity plot of the Simpson index comparing double humanized mice on a mouse chow diet, a high fat diet, or a low fat diet. F) Taxa abundance plot by Order level comparing double humanized mice on a mouse chow diet, a high fat diet, or a low fat diet. G) Taxa abundance plot by Family level comparing double humanized mice on a mouse chow diet, a high fat diet, or a low fat diet.

The introduction of a HFD to the double hu-mice decreased measures of alpha diversity, including the number of unique species per sample or species richness, Simpson’s Diversity Index, and Shannon Diversity Index (Figure 5CDE). We also observed a smaller decrease in species richness in the double hu-mice fed with a LFD compared to the Post-FMT samples from double hu-mice fed with regular mouse chow. However, the HFD fed double hu-mice had the lowest species richness and the double hu-mice fed with a LFD did not have decreased Simpson’s Diversity Index or Shannon Diversity Index compared to the Post-FMT fecal samples from the double hu-mice fed with a regular mouse chow.

When Pre-FMT and human donor fecal sample data was added to the analysis, the Pre-FMT samples had pre-existing low diversity measurements (Supplemental Figure Diet_Comp.pdf). After antibiotic treatment and human FMT (Post-FMT), the double hu-mice on a regular mouse chow diet had increased alpha diversity measurements. After introduction of the HFD, the alpha diversity measurements dropped to near Pre-FMT levels. These data show that dietary fat content plays an important role in regulating gut microbiome diversity. However, the fecal samples from the LFD group also had a decrease in species richness compared to Post-FMT samples, which implicates other factors that may be important for gut microbiome diversity, such as dietary fiber content.

Multiple differences were observed in the composition of the gut microbiome with the three different diets as shown by the heatmaps of bacterial relative abundance for Order and Family taxa levels (Figure 5FG). Significant differences in the relative abundance of gut bacterial taxa between double hu-mice consuming different diets were found using Kruskal-Wallis tests with false discovery rate (FDR) adjusted P values <.05 (Supplemental File Diet_KW.xlsx). Compared to LFD samples, HFD samples had a higher relative abundance of *Verrucomicrobiaceae* (11.*35*%, P=0.0037 FDR) and *Lachnospiraceae* (3.48%, P=0.0025 FDR) and a lower abundance of Firmicutes (−8.71%, P=0.0300 FDR), *Clostridiales* (−5.96%, P=0.0279 FDR), and *Ruminococcaceae* (−2.44%, P=0.0028 FDR).

Compared to Post-FMT samples from double hu-mice fed regular mouse chow, HFD fed double hu-mice had a higher relative abundance of *Streptococcaceae* (17.32%, P=0.0001 FDR), *Bacteroides fragilis* (3.47%, P=0.0001 FDR), *Dorea* (2.64%, P=0.0001 FDR), *Enterobacteriaceae* (1.23%, P=0.0001 FDR), *Enterococcaceae* (0.90%, P=0.0074 FDR), *Desulfovibrionaceae* (0.63%, P=0.0007 FDR) and a lower abundance of *Blautia* (−9.73%, P=0.0001 FDR), *Bacteroidaceae* (−8.30%, P=0.0001 FDR), *Clostridiales* (−5.68%, P=0.0144 FDR), and *Turicibacteraceae* (−5.61%, P=0.0001 FDR).

While our LFD group acted as a calorie matched control for the HFD group, we found the LFD fed double hu-mice also had an altered gut microbial composition compared to Post-FMT samples from double hu-mice fed regular mouse chow. LFD fed double hu-mice had a higher abundance of *Streptococcaceae* (16.16%, P=0.0001 FDR), *Ruminococcaceae* (2.20%, P=0.0102 FDR), *Enterococcaceae* (1.72%, P=0.0001 FDR), and *Proteobacteria* (1.22%, P=0.0443 FDR) and a lower abundance of *Blautia* (−9.97%, P=0.0001 FDR), *Verrucomicrobiaceae* (−8.93%, P= 0.0414*), *Bacteroidaceae* (−7.03%, P=0.0001 FDR), *Turicibacteraceae* (−5.49%, P=0.0001 FDR), and *Erysipelotrichaceae* (−2.47%, P=0.0495*). We found that differences in the both fat and fiber content of the three diets had a large impact on diversity and abundance of the gut microbiome in the double hu-mice model.

A random forest model was trained to predict if the gut microbiome profiles came from double hu-mice that were consuming a HFD or LFD based on the ASV features. The top 15 most important discriminatory features of the model based on area under the ROC curve were then identified. These features were scaled to 100 and plotted along with the average normalized ASV counts for each diet (Supplemental File Diet_Importance.pdf). The top ranked features were ASV107 and ASV476, both from *Oscillospira*, with ASV107 more prevalent with a LFD and ASV476 more prevalent with a HFD. Included in the top 15 features were ASVs from *Bacteroides, Clostridiales, Christensenellaceae, Christensenella, Lachnospiraceae, Clostridium citroniae, Clostridium methylpentosum, Oscillospira* and *Erysipelotrichaceae*. Interestingly, ASV476 from *Oscillospira*, was identified in both the HIV-1 infection and diet random forest models. It was more prevalent in both HIV-1 infected double hu-mice and in double hu-mice consuming a HFD. Using a HFD we successfully induced microbial dysbiosis in our double hu-mice model. We found that compared to regular mouse chow, a diet consisting of high-fat content and a lack of fiber significantly changed the gut microbiome composition, including a decrease in alpha diversity.

### High-fat diet induced gut microbial changes were associated with increased systemic inflammation and immune activation

To evaluate if the HFD fed double hu-mice had elevated levels of systemic inflammation, we measured the levels of inflammatory cytokines in plasma using multiplex immunoassays. Double hu-mice consuming a HFD had significantly higher levels of IL-1β than mice on the LFD (Figure 6A). Interestingly, the levels of IL-1β increased in the HFD fed double hu-mice in each timepoint tested (Figure 6B). The levels of inflammatory cytokines IL-6 and IFN-γ were both significantly higher in mice consuming the HFD compared to the LFD (Figure 6CDEF). However, levels of TNF-α were not significantly different between the two groups of mice, with the highest measured level found in the LFD group at 0.5 weeks post diet initiation (Figure 6GH). Feeding with the HFD quickly raised the levels of systemic inflammatory markers IL-1β, IL-6, and IFN-γ, with progressively increased IL-1β at each measured timepoint. It was clear that the HFD not only led to microbial dysbiosis, but also increased the levels of systemic inflammation.

**Figure 6.**
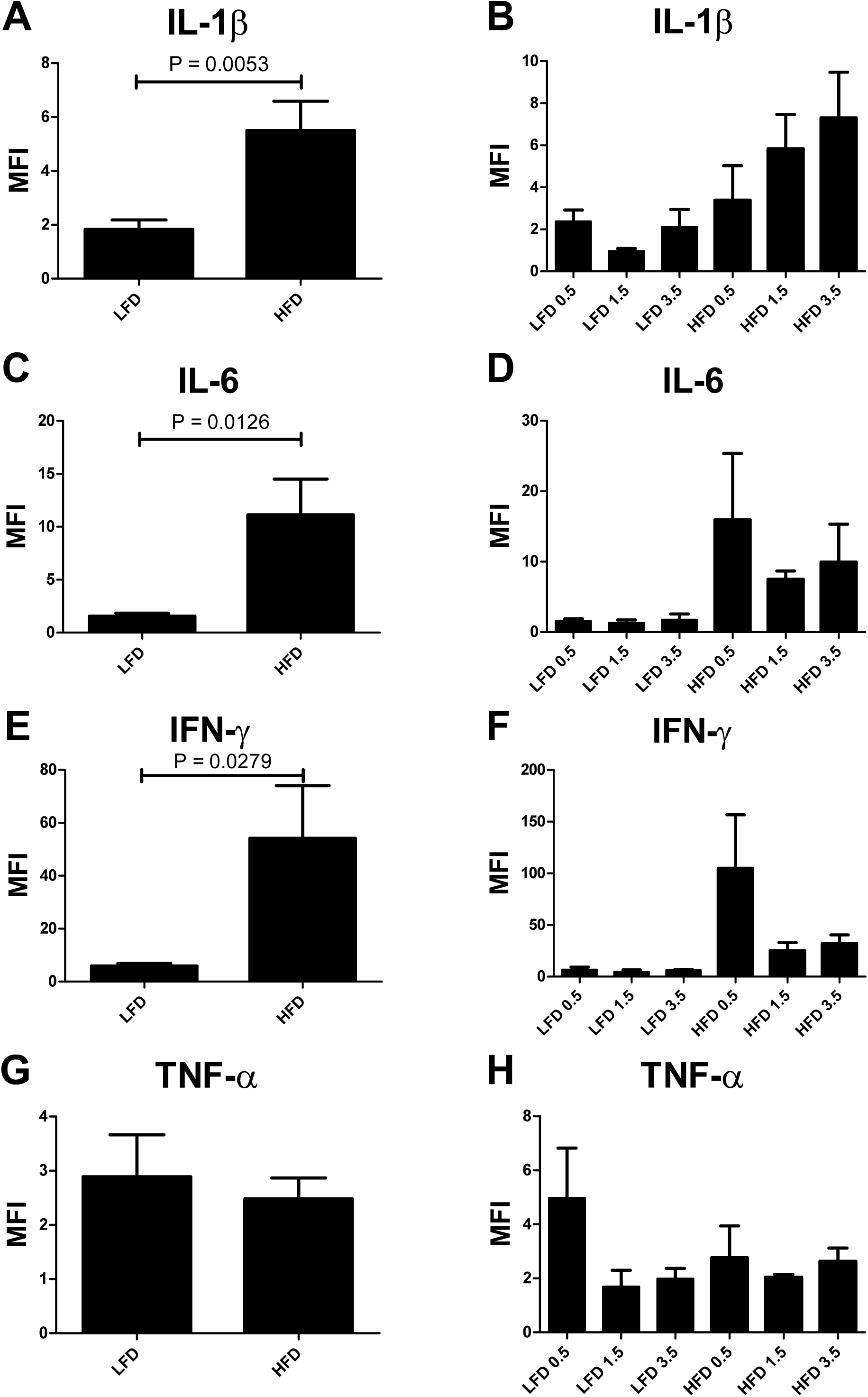
Double hu-mice fed a high fat diet had increased systemic human inflammatory cytokines. Human inflammatory cytokine measures from plasma of double humanized mice. Samples were collected from double humanized mice 0.5, 1.5, and 3.5 weeks post low fat diet (LFD) or high fat diet (HFD) initiation. Cytokine levels shown by mean fluorescence intensity (MFI). A) All samples IL-1β B) Longitudinal IL-1β C) All samples IL-6 D) Longitudinal IL-6 E) All samples IFN-γ F) Longitudinal IFN-γ G) All samples TNF-α H) Longitudinal TNF-α.

Along with inflammation, increased immune activation is an important pathogenic factor for enhancing HIV-1 transmission and pathogenesis. Using flow cytometry, we measured immune activation of human immune cells in peripheral blood (Figure 7AB). Unlike the HIV-1 infected double hu-mice, CD4+ T cell populations in both HFD and LFD groups (Table 1) remained steady. The activated T cell populations of CD4+ CD38+ and CD8+ CD38+ were increased in the HFD group compared to the LFD group. Additionally, the CD4+ HLA-DR+ population increased over time in the HFD fed group and the largest population of CD8+ HLA-DR+ cells were observed at 3 weeks after diet initiation in the HFD group. The expression of CD69 did not change with the introduction of the HFD or LFD in either CD4+ or CD8+ T cells. The HFD significantly altered the gut microbial composition in double hu-mice and was associated with both increased systemic inflammation and immune activation.

**Figure 7.**
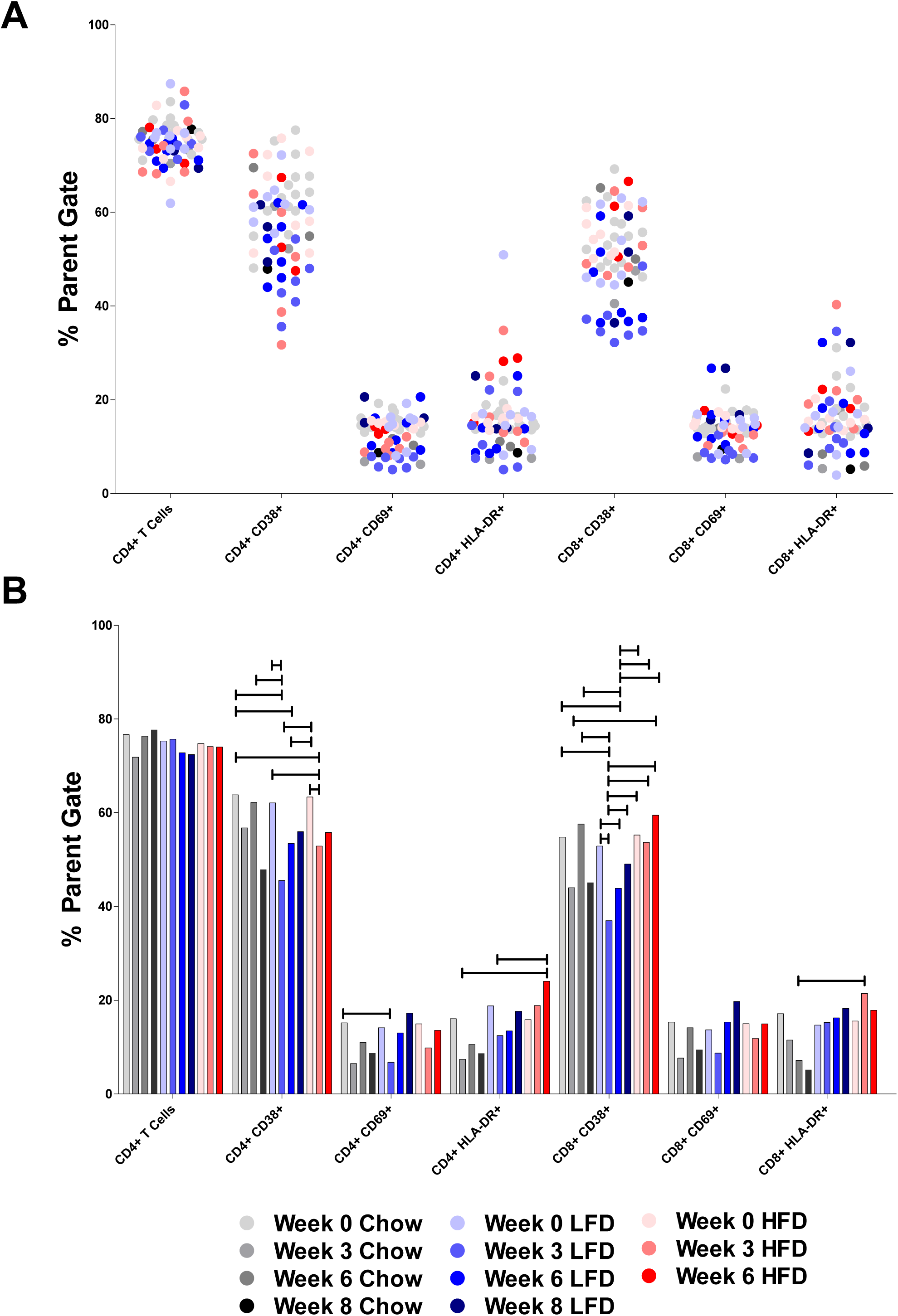
Double hu-mice fed a high fat diet had increased systemic human immune cell activation. A) Human immune cell populations from peripheral blood of double humanized mice up to 8 weeks on a regular mouse chow (Chow), low fat (LFD), or high fat diet (HFD). All immune populations were lymphocyte+, human CD45+ and mouse CD45-, and human CD3+) and are represented by the percentage of their parent gate. B) Percentage of peripheral blood human immune cell populations shown as a mean for the longitudinally collected double humanized mice on a chow, LFD, or HFD. For each population of human immune cells a multiple comparison test for significance was performed between the sample groups (ANOVA with Tukey Test with adjusted P values < 0.05).

### Relationships between the gut microbiome and systemic inflammation and immune activation

To better understand the bidirectional relationship between the gut microbiota and immune system, we compared plasma derived inflammatory cytokine levels of IL-1β, IL-6, IFN-γ, and TNF-α with matched gut microbiome profiles from our double hu-mice experiments (Supplemental File Correlations.xlsx). Ten ASVs were significantly correlated with IL-1β, including 7 from *Clostridiales* and 3 from *Klebsiella* (Supplemental File Cytokine_Correlations.pdf). None of the significant ASVs were found in the sequenced human donor samples. Interestingly, 9 of 10 ASVs correlated with IL-1β were also significantly correlated with IL-6. Additionally, IL-6 was significantly correlated with ASV726 *Christensenellaceae*, which was also found in human donor samples. 20 ASVs were significantly correlated with IFN-γ, of which 8 were also found in human donor samples. Significantly correlated ASVs included *Bacteroides eggerthii, Blautia obeum*, and *Coprococcus catus*. Several ASVs mapped to *Clostridiales*, including two from *Oscillospira*. Twenty-three ASVs were significantly correlated with TNF-α, 10 of which were also found in human donor samples. Some of the significant ASVs were mapped to *Coriobacteriaceae, Bacteroides uniformis, Rikenellaceae, Enterococcus, Blautia, Oscillospira; Citrobacter*, and *Klebsiella*. Interestingly, many the ASVs that correlated with IL-1β and IL-6 were the same and these ASVs were not found in the human donor samples. However, ASVs that correlated with IFN-γ, and TNF-α were much more likely to be found in human donor samples. While not a direct sign of causation, many of the identified ASVs came from bacteria that have established interactions with the immune system or are known to be potentially pathogenic.

To better understand the relationship between the human-like gut microbiome and human immune cell activation, we compared flow cytometry data derived from peripheral blood cells with matched gut microbiome profiles from our double hu-mice experiments (Supplemental File Correlations.xlsx). We identified 54 significant correlations corresponding to 37 unique ASVs, including 6 ASVs that can be found in the human donor samples. ASV14 *Bacteroides* and ASV242 *Clostridiales*, were positively correlated with several markers of CD8+ T cell immune activation (Supplemental Figure Immune_Correlations.pdf). ASV280 *Barnesiellaceae* was negatively correlated with CD4+ T cells and positively correlated with activation of CD8 T cells. Interestingly, ASV280 was also positively correlated with the level of plasma IFN-γ. Additionally, several more ASVs were negatively correlated with CD4 T cells including ASV535 *Christensenella*, ASV606 *Christensenellaceae*, and ASV644 *Ruminococcaceae*. ASV717 *Clostridiales* was negatively correlated with human CD45+ immune cells and positively correlated with CD8 T cell immune activation, while ASV870 *Clostridiales* was also positively correlated with CD8 T cell immune activation. ASV955 *Oscillospira* and ASV965 *Oscillospira* were both negatively correlated with CD4 T cells and positively correlated with CD8 T cell activation. Additionally, several ASVs were identified to be positively correlated with both markers for CD8 T cell activation and plasma IFN-γ. These ASVs included members of *Ruminococcaceae, Lachnospiraceae*, and *Bacteroides eggerthii*. We were able to identify several ASVs that correlated with both CD4 T cell loss and CD8 T cell activation. Further, ASVs that correlated with IFN-γ were often also correlated with CD8 T cell activation.

## Discussion

The gut microbiome and immune system have a complex and interdependent relationship. As we previously reported and also confirmed in this study, we found that the murine gut microbiome of regular hu-BLT mice (hu-mice) had lower levels of diversity and differed greatly in composition from the microbiomes of our human fecal donors [58, 59]. These large differences between the gut microbiomes of murine origin from hu-mice and humans may limit the translatability of experimental results [55, 56]. In this study, the double hu-mice harboring both a human immune system and human gut microbiome allowed for the study of the relationship between the human gut microbiome and human immune system during HIV-1infection and a HFD. We tracked the compositional changes to the gut microbiome before and after human FMT, during HIV-1 infection for up to 12 weeks, and HFD for up to 7 weeks. During these longitudinal studies, we also measured plasma pro-inflammatory cytokines and quantified immune activation of human CD4 and CD8 T cells isolated from peripheral blood and spleen.

HIV-1 infection profoundly alters the human immune system with long lasting consequences, such as persistent immune activation and inflammation despite suppressive ART [14, 24, 70, 71]. The gut and gut microbiome may play an important role in many aspects of HIV-1 mucosal transmission, CD4+ T cell death, and the elevated risk of comobidities in PLWH on ART. Therefore, changes in the gut microbiome during HIV-1 infection have been widely studied [19, 20, 72-92]. However, many of these human studies varied in sampling and analytical methods, as well as geography, age, sex, diet, and lifestyle choices of the study subjects. Further, it is difficult to study very early stages of HIV-1 infection and there is often a wide range of disease progression rate, timing of ART treatment, and treatment outcomes in human studies. As such, it has been difficult to discern changes in the gut microbiome due to HIV-1 infection or other factors across the various studies. One major finding in earlier studies was that the gut microbiome profiles of HIV infection had higher levels of *Prevotella [72, 77, 81, 86]*. However, studies controlling for lifestyle choices, such as men who have sex with men (MSM), have not found significant changes in *Prevotella* levels due to infection, but rather have found high levels of *Prevotella* in MSM [19, 80, 87, 89, 93-96]. While important questions remain as to the impact of gut microbiome profiles with a high abundance of *Prevotella* on HIV-1 transmission, pathogenesis, and treatment, this is a clear example of the difficulty of studying the gut microbiome in humans due to the many confounding factors.

Non-human primates (NHP) infected with SIV as animal models have successfully been used to study many aspects of HIV-1 pathogenesis. However, the subtle changes in the gut microbiome of SIV infected NHP were not consistent with the more significant changes observed in HIV-1 infected human studies [97-99]. In this study, we utilized a unique double hu-mice model to complement the studies performed in humans and NHP. Compared to NHP, our double hu-mice model has the advantage of using HIV-1 instead of SIV for infection and have both a human-like gut microbiome and human immune system. Importantly, we can use the model to track changes in the gut microbiome longitudinally, from very early to chronic disease stages, while controlling for many of the confounding factors that make studies in humans difficult.

In this study, compared to uninfected double hu-mice, infected double hu-mice had a higher relative abundance of *Bifidobacteriaceae* and *Ruminococcaceae* and a lower abundance of *Lactobacillaceae* and *Turicibacteraceae*. Compared to pre-infection samples, infected double hu-mice had a higher relative abundance of *Bacteroidaceae, Bifidobacteriaceae, Clostridiaceae, Rikenellaceae*, and *Ruminococcaceae* and a lower relative abundance of *Erysipelotrichaceae, Lachnospiraceae, and Verricomicrobiaceae*. A meta-analysis of sex and lifestyle matched controlled studies found that HIV-infected populations were enriched with *Erysipelotrichaceae, Enterobacteriaceae, Desulfovibrionaceae*, and *Fusobacteria* and depleted of *Lachnospiraceae, Ruminococceae, Bacteroides*, and *Rikenellaceae[93]*. In this study, we did not find large shifts in the gut microbiome due to HIV-1 infection alone. We believe that housing the hu-mice in controlled environments without natural exposure to outside pathogens may account for why we did not observe increases in relative abundance of bacteria like *Erysipelotrichaceae, Enterobacteriaceae, Desulfovibrionaceae*. Further, increases in *Fusobacteria* may be linked to ART treatment itself and not untreated HIV-1 infection. The double hu-mice model may be further improved upon by potentially introducing outside microbes during infection and by studying the impact of ART use by itself and during treatment of HIV-1 infection.

The altered gut microbiome due to HIV-1 infection may also play an important role in HIV-1 pathogenesis, including the loss of CD4 T cells, inflammation, and immune activation. In this study, we found that HIV-1 infection of double hu-mice increased the levels of systemic pro-inflammatory cytokines IL-1β, IL-6, IFN-γ, and TNF-α. As expected, HIV-1 infected double hu-mice had significantly decreased CD4 T cells and increased immune activation manifested in three populations of activated CD8 T cells (CD38+, CD69+, or HLA-DR+). Moreover, CD4 T cell loss and T cell immune activation was also confirmed in lymphocytes isolated from the spleens of infected double hu-mice. We also investigated the relationships between the observed immunopathogenesis with the gut microbiome. We found nine ASVs that were negatively correlated with CD4 T cells including ASVs from *Barnesiellaceae, Christensenellaceae, Lachnospiraceae, Oscillospira*, and, *Ruminococcus*. ASVs from *Bacteroidales* and *Odoribacter* were correlated with increased CD4+ CD38+ populations. Thirty-seven ASVs positively correlated with increased CD8 T cell activation, of which 8 were also positively correlated with increased levels of plasma IFN-γ. These 8 ASVs included members of B*arnesiellaceae, Lachnospiraceae, Ruminococcaceae, Oscillospira*, and *Bacteroides eggerthii*. Our random forest model trained to distinguish gut microbiome profiles of HIV-1 infected and uninfected double hu-mice identified several bacteria with known links to the immune system including ASVs from *Ruminococcus, Lachnospiraceae, Christensenellaceae, Oscillospira, Dorea, Butyricicoccus pullicaecorum and Blautia producta*. While not a direct measure of causation, these correlations provide a foundation for future study in order to narrow down key groups of bacteria that play a role in immunopathogenesis during HIV-1 infection.

Another important aspect of this study was the establishment of gut microbial dysbiosis in double hu-mice using a HFD. Establishing a state of microbial dysbiosis from a gut microbiome engrafted from a healthy human fecal donor sample was an important first step in order to determine the role the gut microbiome in HIV-1 rectal transmission. We showed that a HFD quickly lowered the alpha diversity and changed the composition of the gut microbiome. We also found that double hu-mice that consumed a HFD had increased levels of pro-inflammatory cytokines IL-1β, IL-6, IFN-γ, along with increased populations of activated CD4 and CD8 T cells. We showed that CD4+ CD38+ population, which were decreased as a result of HIV-1 infection, were increased with a HFD. In our study, we showed that diet and the corresponding gut microbial dysbiosis can have drastic systemic effects on inflammation and immune activation. Future studies are needed to determine if gut microbial dysbiosis impacts susceptibility to HIV-1 rectal transmission and subsequent pathogenesis.

We also would like to point out the limitations of our study. First, we only characterized the gut bacteria and did not investigate the gut virome. During HIV-1 infection there is an expansion of the virome which may contribute to the observed persistent immune activation and inflammation in PLWH [80]. Second, metagenomic sequencing would also capture functional changes in the gut microbiota important to HIV-1 pathogenesis. Last, the double hu-mice model could be expanded to include patient derived fecal donor samples and fecal donors with diverse gut microbiome profiles. Going forward, we believe that double hu-mice could provide a complimentary model to help answer some of the outstanding questions about the relationship between the gut microbiome and HIV-1 infection.

## Conclusions

Here, we describe the changes in the gut microbiome and human immune system due to HIV-1 infection and a HFD using our double hu-mice model. HIV-1 infection led to changes in the composition of the human-like gut microbiome that was associated with CD4 T cell loss and high levels of inflammation and immune activation. Microbial dysbiosis was quickly established in double hu-mice through feeding a HFD and led to systemic immune activation and inflammation. We also identified a subset of gut bacteria that was closely associated with systemic inflammation and immune activation in double hu-mice infected with HIV-1 or fed a HFD. Importantly, this study demonstrated how the double hu-mice model can be used to longitudinally study the complex in vivo interactions of the gut microbiome and human immune system.

## Methods

### Generation of hu-BLT mice

All methods described here were conducted as we previously reported in accordance with Institutional Animal Care and Research Committee (IACUC)-approved protocols at the University of Nebraska-Lincoln (UNL)[57, 60, 61, 64]. The IACUC at the University of Nebraska-Lincoln (UNL) has approved two protocols related to generating and using hu-BLT mice, including Double Hu-Mice. Additionally, the Scientific Research Oversight Committee (SROC) at UNL has also approved the use of human embryonic stem cells and fetal tissues, which are procured from the Advanced Bioscience Resources for humanized mice studies (SROC# 2016—1-002).

Briefly, 6-to 8-week-old female NSG mice (NOD.*Cg-Prkdc*^*scid*^*Il2rg*^*tm1Wjl*^/SzJ, catalog number 005557; (Jackson Laboratory) were housed and maintained in individual microisolator cages in a rack system capable of managing air exchange with prefilters and HEPA filters. Room temperature, humidity, and pressure were controlled, and air was also filtered. Mice given autoclaved and acidified drinking water ab libitum and were fed one of the following diets determined by the experimental group, irradiated Teklad global 14% protein rodent chow (Teklad 2914), irradiated Teklad Rodent Diet With 10 kcal% Fat (No Sucrose) (Teklad K12450Ki), or irradiated Teklad Rodent Diet With 60% kcal% Fat (Teklad D12492i). On the day of surgery, mice received whole-body irradiation at the dose of 12 cGy/gram of body weight with the RS200 X-ray irradiator (RAD Source Technologies, Inc., GA). Each irradiated mouse was given 130-170 ul of a mixture of Ketamine/Xylazine (0.27 ml of Ketamine at the concentration of 100 mg/ml and 0.03 ml of Xylazine at the concentration of 100 mg/ml to 2.7 ml of sterile saline) by intraperitoneal (IP) injection for anesthesia. Additionally, each mouse was given 100 ul Buprenex (half-life 72 hours, 1mg/kg of body weight) by subcutaneous injection for long lasting pain management and 100 ul (858 ug) Cefazolin by IP injection for antibiotic prophylaxis. Isofluorane gas at 3-5% was given if additional anesthesia was needed at any point during surgery. After proper levels of anesthesia were verified by pedal reflex testing, each mouse was implanted with one piece of human fetal thymic tissue fragment sandwiched between two pieces of human fetal liver tissue fragments within the murine left renal capsule. Within 6 hours of surgery, mice were injected via the tail vein with 1.5 × 10^5^ to 5 × 10^5^ CD34^+^ hematopoietic stem cells isolated from human fetal liver tissues. Human fetal liver and thymus tissues were procured from Advanced Bioscience Resources (Alameda, CA). After 10 weeks, human immune cell reconstitution in peripheral blood was measured by a fluorescence-activated cell sorter (FACS) Aria II flow cytometer (BD Biosciences, San Jose, CA) using antibodies against mCD45-APC, hCD45-FITC, hCD3-PE, hCD19-PE/Cy5, hCD4-Alexa 700, and hCD8-APC-Cy7 (catalog numbers 103111, 304006, 300408, 302209, 300526, and 301016, respectively; BioLegend, San Diego, CA). Raw data were analyzed with FlowJo (version 10.0; FlowJo LLC, Ashland, OR). All humanized mice used in this study had high levels of human immune cell reconstitution with an average of 89.4% hCD45+ cells in peripheral blood 10 weeks post-surgery. The mice were randomly assigned into experimental groups with similar immune reconstitution levels. Mice were euthanized at humane study endpoints with carbon dioxide followed by cervical dislocation in accordance with approved Institutional Animal Care and Research Committee (IACUC)-approved protocols at the University of Nebraska-Lincoln (UNL). Following the approved protocols, animals were euthanized before or at the point of observed impaired ambulation, prolonged drowsiness or aversion to activity, lack or physical or mental alertness, prolonged inappetence, difficulty breathing, chronic diarrhea or constipation, inability to remain upright, or at the discretion of the Veterinary Staff.

### Antibiotic treatment

A broad-spectrum antibiotic cocktail was prepared fresh daily consisting of Metronidazole (1 g/L), Neomycin (1 g/L), Vancomycin (0.5 g/L), and Ampicillin (1 g/L). The antibiotic cocktail was given to the mice ad libitum in the drinking water along with grape flavored Kool-Aid to improve palatability. Control group mice were given only grape flavored Kool-Aid in the drinking water. During antibiotic treatment, cages were changed daily to limit re-inoculation of pre-existing bacteria to the mice due to their coprophagic behavior. Antibiotics were given for 14 days for all double hu-mice. Post-antibiotic treatment, mice were given autoclaved non-acidified deionized drinking water. Body weight was carefully monitored during this time and If needed, mice were treated with Intraperitoneal (IP) injections of Ringer’s solution to mitigate any effects of dehydration.

### Donor samples and Fecal transplant

At 24 and 48 hours after the completion of antibiotic pre-treatment, mice were given 200 ul of human fecal material via oral gavage. OpenBiome supplied 3 FMT Upper Delivery Microbiota Preparations from 3 different healthy human donors (Donor 65, Donor 74, Donor 82). Samples were thawed once before fecal transplant to aliquot the samples within an anaerobic chamber. During this step, an equal portion of each of the samples were mixed together to create an unbiased human donor sample.

### HIV-1 infection and q-RT-PCR

To infect double hu-mice with HIV-1, mice were intraperitoneally injected with 4.5*10^5 TCID of an equal mixture of HIV-1_SUMA_ and HIV-1_JRCSF_. To verify infection, plasma viral RNA was extracted using a QIAamp ViralRNA minikit (Qiagen). Plasma viral load was conducted using reverse transcriptase quantitative PCR (qRT-PCR) on a C1000 ThermalCycler and the CFX96 Real-Time system (Bio-Rad) and the TaqMan FastVirus 1-Step master mix (Life Technologies). As previously reported, the following primers were used for the plasma viral load assay: Forward Primer: GCCTCAATAAAGCTTGCCTTGA; Reverse Primer: GGGCGCCACTGCTAGAGA; Probe: /56-FAM/CCAGAGTCA/ZEN/CACAACAGACGGGCACA/3IABkFQ/[64].

### Multiplex immunoassay for measuring plasma cytokines

Plasma from double hu-mice was tested for the following inflammatory cytokines: IFN-γ, IL-1β, IL-2, IL-4, IL-6, IL-10, IL-12 p70, IL-17A, TNFα using the ProcartaPlex high sensitivity 9-Plex Human Panel (EPXS090-12199-901, Thermofisher Scientific, Waltham, MA). Samples were measured using a Luminex MAGPIX instrument (Luminex Corporation, Austin, TX).

### Lymphocytes isolation and immune activation flow cytometry panel

Spleens from euthanized double hu-mice were placed on a strainer with 70-μm nylon mesh (Cat#22363548, Fisher Scientific) and the strainer was placed on a 50ml Centrifuge tube (Corning). Spleen tissue was gently pressed with the flat end of a 5-ml syringe to release the splenocytes into cell cultural medium that contained 90% RPMI-1640 (Cat#11875, Life technologies) supplemented with 10% heat-inactivated fetal bovine serum [HI-FBS] (Cat#SH30071.03, Thermo Scientific), penicillin [100 IU/ml]-streptomycin [100 ug/ml] (Quality Biological, Inc), 2mM/ml L-glutamine (Cat#25-005-CI, Corning). Slowly, the splenocyte suspension was layered onto Histopaque-1077 (Sigma-Aldrich), and centrifuge at 350×g for 30 mins at room temperature. The “buffy coat” mononuclear cells layer were transferred into a 50ml Centrifuge tube (Corning) and washed with cold PBS. Human immune cell activation in peripheral blood and lymphocytes isolated from the spleen were measured by a fluorescence-activated cell sorter (FACS) Aria II flow cytometer (BD Biosciences, San Jose, CA) using antibodies against mCD45-APC, hCD45-FITC, hCD3-PE, hCD4-Alexa Fluor 700, hHLA-DR-BV421, hCD38-PE-Cy5, hCD69-BV785, hCD8a-APC-Cy7, mCD45-APC, Viability-APC (catalog numbers 304006, 300408, 300526, 307636, 303508, 310932, 301016, 103112 (BioLegend, San Diego, CA), and 65-0864-14 (eBioscience, San Diego, CA). Raw data were analyzed with FlowJo (version 10.0; FlowJo LLC, Ashland, OR).

### Mouse fecal collection and DNA extraction

Individual mice were placed into autoclaved paper bags within a biosafety hood until fresh fecal samples were produced. Fecal samples were stored in 1.5 ml Eppendorf tubes at −80 °C until DNA extraction. DNA was extracted from the fecal samples using the phenol:chloroform:isoamyl alcohol with bead beating method described previously [100]. Briefly, fecal samples were washed three times with 1 ml PBS buffer (pH 7). After the addition of 750 ul of lysis buffer, samples were transferred to tubes containing 300 mg of autoclaved 0.1 mm zirconia/silica beads (Biospec). 85 ul of 10% SDS solution and 40 ul of Proteinase K (15mg/ml, MC500B Promega) were added and samples were incubated for 30 minutes at 60° C. 500 ul of Phenol:Chloroform:Isoamyl alcohol (25:24:1) was added and then samples were vortexed. Samples were then put into a bead beater (Mini-beadbeater 16 Biospec) for 2 minutes to physically lyse the cells. The upper phase of the sample was collected and an additional 500ul of Phenol:Chloroform:Isoamyl alcohol (25:24:1) was added. After samples were vortexed and spun down, the DNA in the upper phase was further purified twice with 500 ul of Phenol:Chloroform:Isoamyl alcohol (25:24:1). and was then precipitated with 100% Ethanol (2.5 x volume of sample) and 3M Sodium acetate (.1 x volume of sample) overnight at −20° C. Samples are then centrifuged and dried at room temperature. DNA was resuspended in 100 ul of Tris-Buffer (10mM, pH8) and stored at −20° C. DNA samples were quality checked by nanodrop (ND-1000 Nanodrop).

### 16S rRNA gene sequencing

16S rRNA gene sequencing was performed at the University of Nebraska Medical Center Genomics Core Facility. DNA normalization and library prep were performed followed by V3-V4 16S rRNA amplicon gene sequencing using a MiSeqV2 (Illumina) The following primer sequences were used: (Primer sequences: Forward Primer = 5’ TCGTCGGCAGCGTCAGATGTGTATAAGAGACAGCCTACGGGNGGCWGCAG 16S Amplicon PCR Reverse Primer = 5’ GTCTCGTGGGCTCGGAGATGTGTATAAGAGACAGGACTACHVGGGTATCTAATCC Illumina overhangs: Forward overhang: 5’ TCGTCGGCAGCGTCAGATGTGTATAAGAGACAG□[locusspecific sequence] Reverse overhang: 5’ GTCTCGTGGGCTCGGAGATGTGTATAAGAGACAG□[locusspecific sequence]).

### Generation of the amplicon sequence variant table and data analysis

Illumina-sequenced paired-end fastq files were demultiplexed by sample and barcodes were removed by the sequencing facility. The University of Nebraska Holland Computer Center Crane cluster was used to run the DADA2 v1.8 R package in order to generate an amplicon sequence variant (ASV) table[101]. The DADA2 pipeline was performed as follows, sequences were filtered and trimmed during which any remaining primers, adapters, or linkers were also removed. The sequencing error rates were estimated using a random subset of the data. Dereplication of the data combined all identical sequencing reads into unique sequences with a corresponding abundance. The core sample inference algorithm was then applied to the dereplicated data. The forward and reverse reads were then joined to create the full denoised sequences and an initial ASV table was generated. Any sequences outside the expected length for the V3-V4 amplicon were then filtered from the table. Chimeric sequences were then removed and a final ASV table was generated. Taxonomy was assigned using the Greengenes 13.8 database and RDP Classifier with a minimal confidence score of 0.80 [102, 103]. Analysis was performed using R package mctoolsr and samples were rarified to 4630 ASVs for downstream analysis. GraphPad Prism 5 and Tableau were used to create some figures. Correlations were performed in R using the rcorr.adjust function in the Hmisc package to compute matrices Spearman correlations along with the pairwise p-values among the correlations. The p-values were corrected for multiple inference using Holm’s method. The random forest models and accompanied variable importance values were generated in R using the randomForest package.

## Supporting information

Supplemental Figure HIV_HuMice

Supplemental File HIV_HuMice_KW

Supplemental File HIV_KW

Supplemental Figure HIV_Composition

Supplemental Figure HIV_Importance

Supplemental Figure HIV_Spleen

Supplemental Figure Diet_Comp

Supplemental File Diet_KW

Supplemental Figure Diet_Importance

Supplemental File Correlations

Supplemental Figure Cytokine_Correlations

Supplemental Figure Immune_Correlations

## Declarations

### Consent for publication

Not applicable

### Availability of data and material

The datasets generated during the current study are available in the NCBI SRA repository, [https://www.ncbi.nlm.nih.gov/bioproject/PRJNA612824].

### Authors’ contributions

LD and QL designed the experiments and wrote the manuscript. LD performed experiments and analyzed the data. ART provided input on experimental design and manuscript preparation.

## Acknowledgments

We would like to thank Pallabi Kundu, Rachel Kubik, Saroj Lohani, Yilun Chang, and Jianshiu Zhang for their assistance in generating hu-BLT mice. We would like to acknowledge the UNMC Genomics Core Facility who receives partial support from the Nebraska Research Network In Functional Genomics NE-INBRE P20GM103427-14, The Molecular Biology of Neurosensory Systems CoBRE P30GM110768, The Fred & Pamela Buffett Cancer Center - P30CA036727, The Center for Root and Rhizobiome Innovation (CRRI) 36-5150-2085-20, and the Nebraska Research Initiative. We would like to thank University of Nebraska—Lincoln Life Sciences Annex and their staff for their assistance.

