## Supplementary figures and images for "Alterations in the gut microbiome by HIV-1 infection or a high-fat diet associates with systemic immune activation and inflammation in double humanized-BLT mice"

### Supplemental Figure Cytokine_Correlations

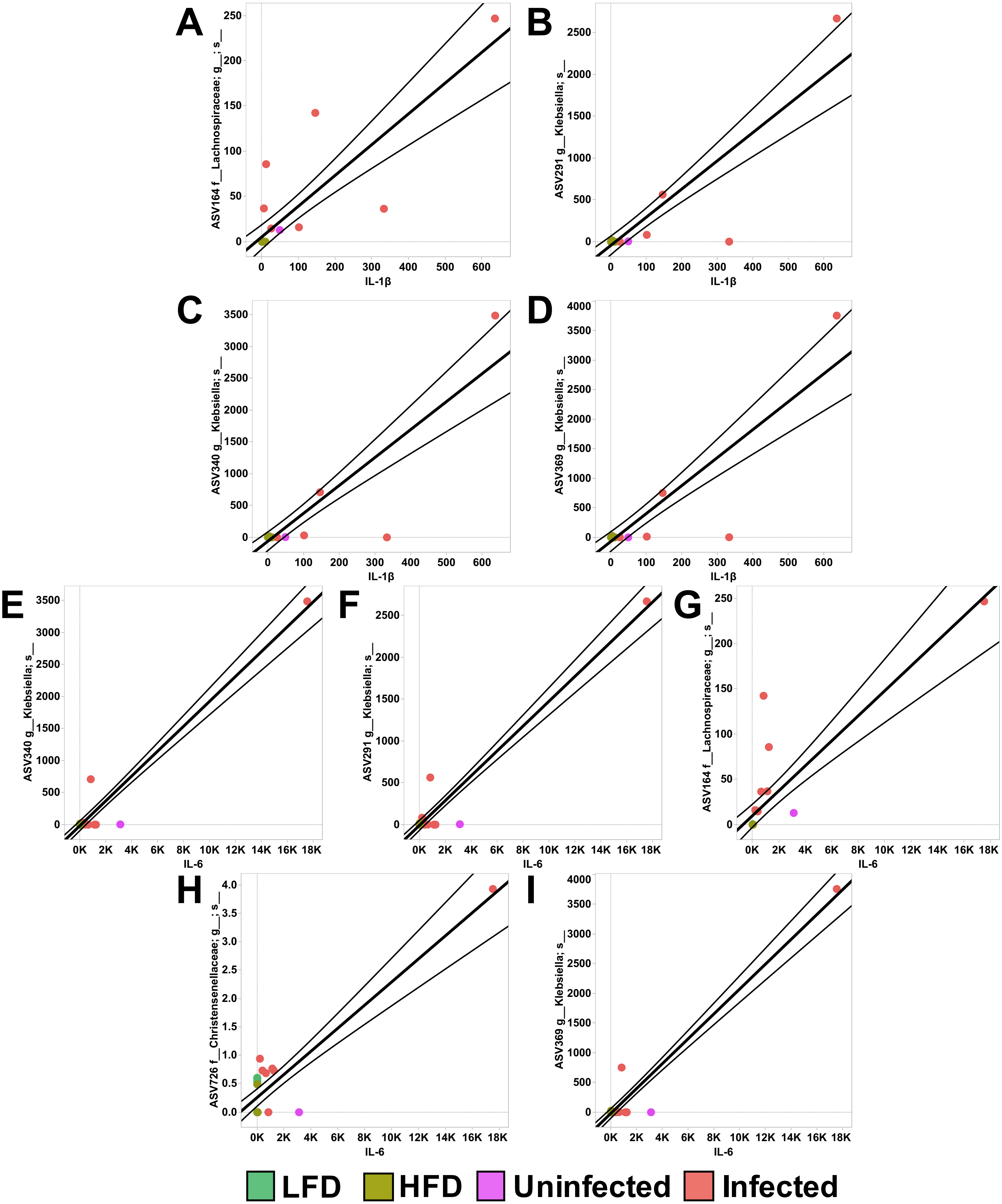

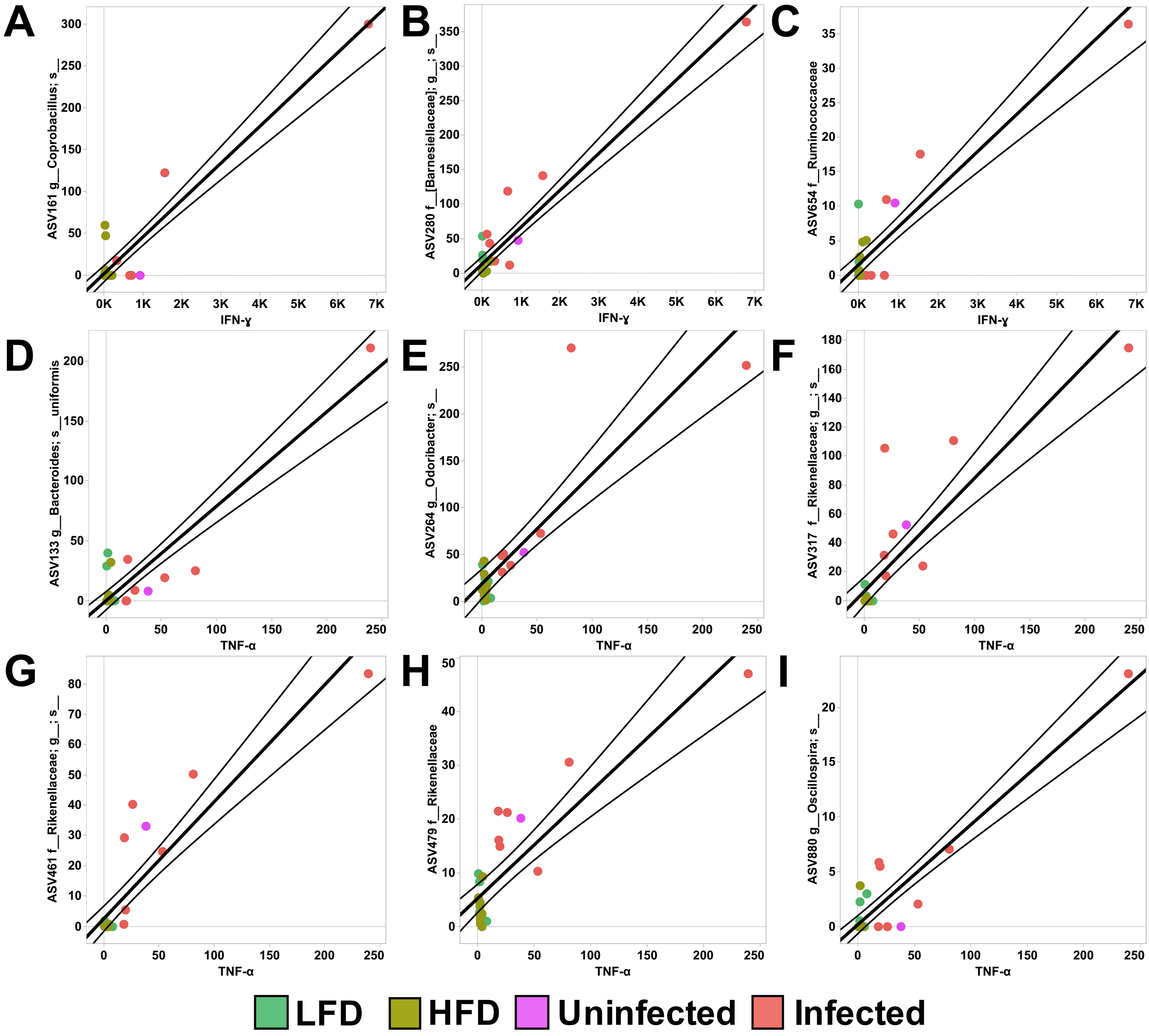

### Supplemental Figure Diet_Comp

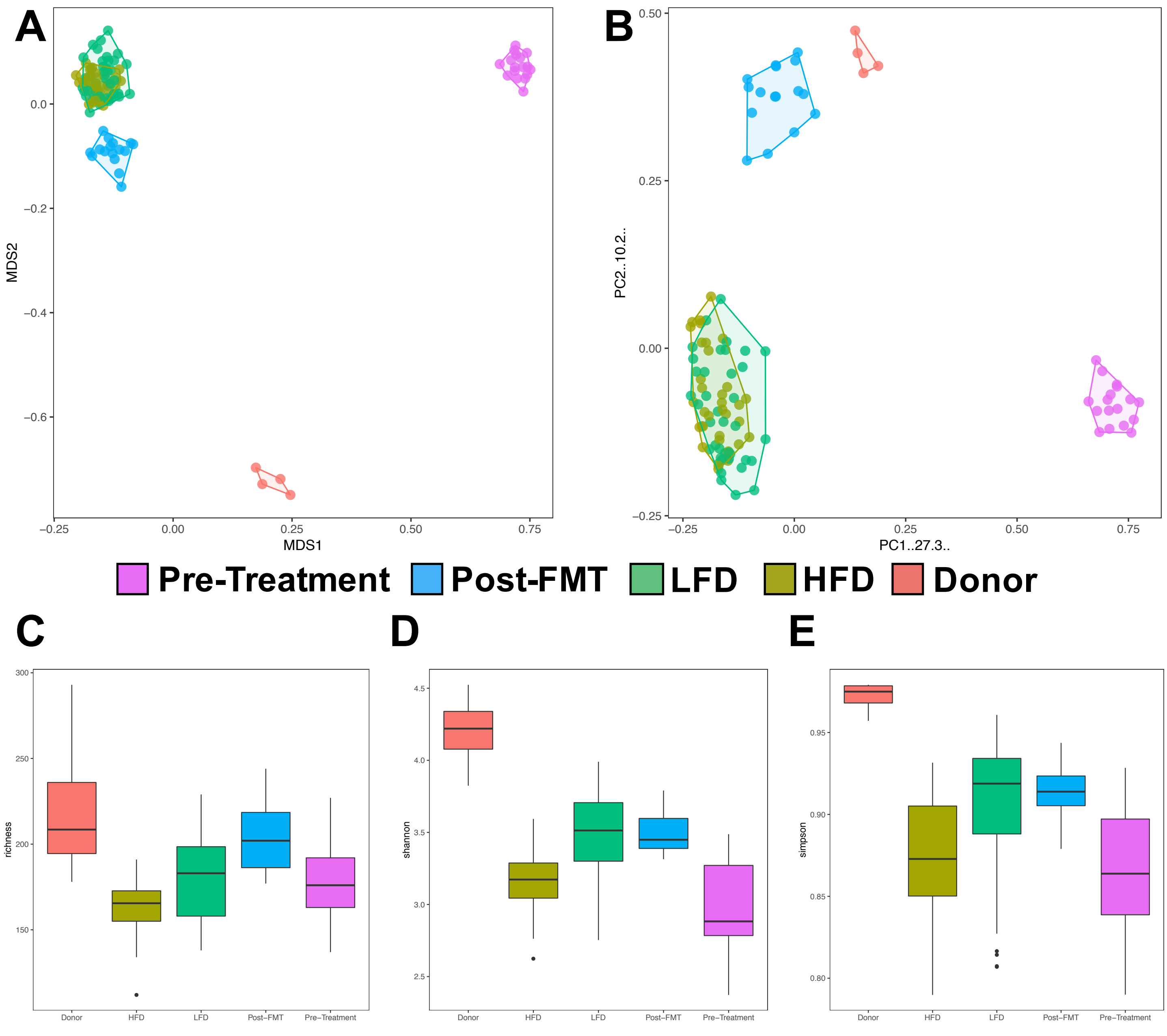

### Supplemental Figure Diet_Importance

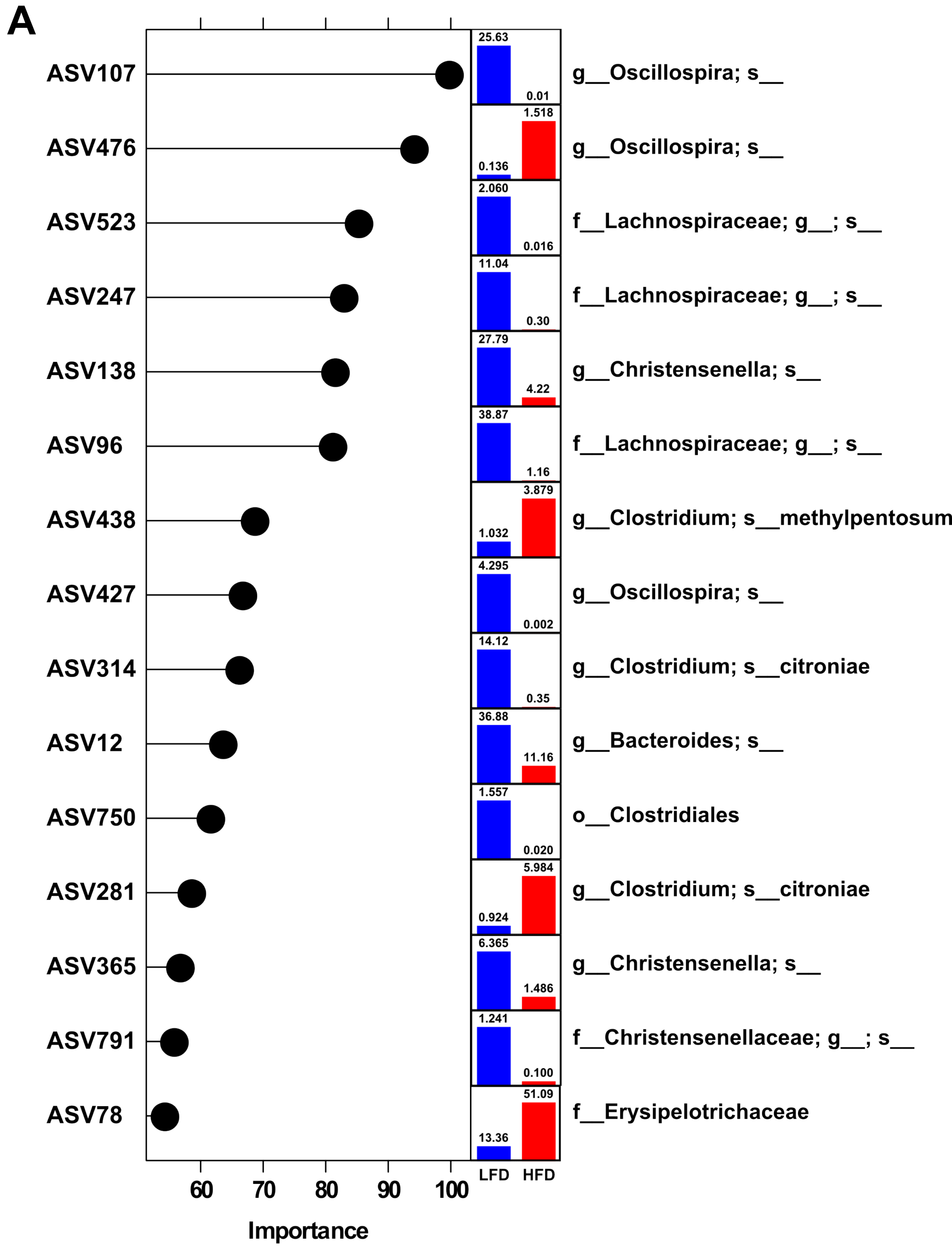

### Supplemental Figure HIV_HuMice

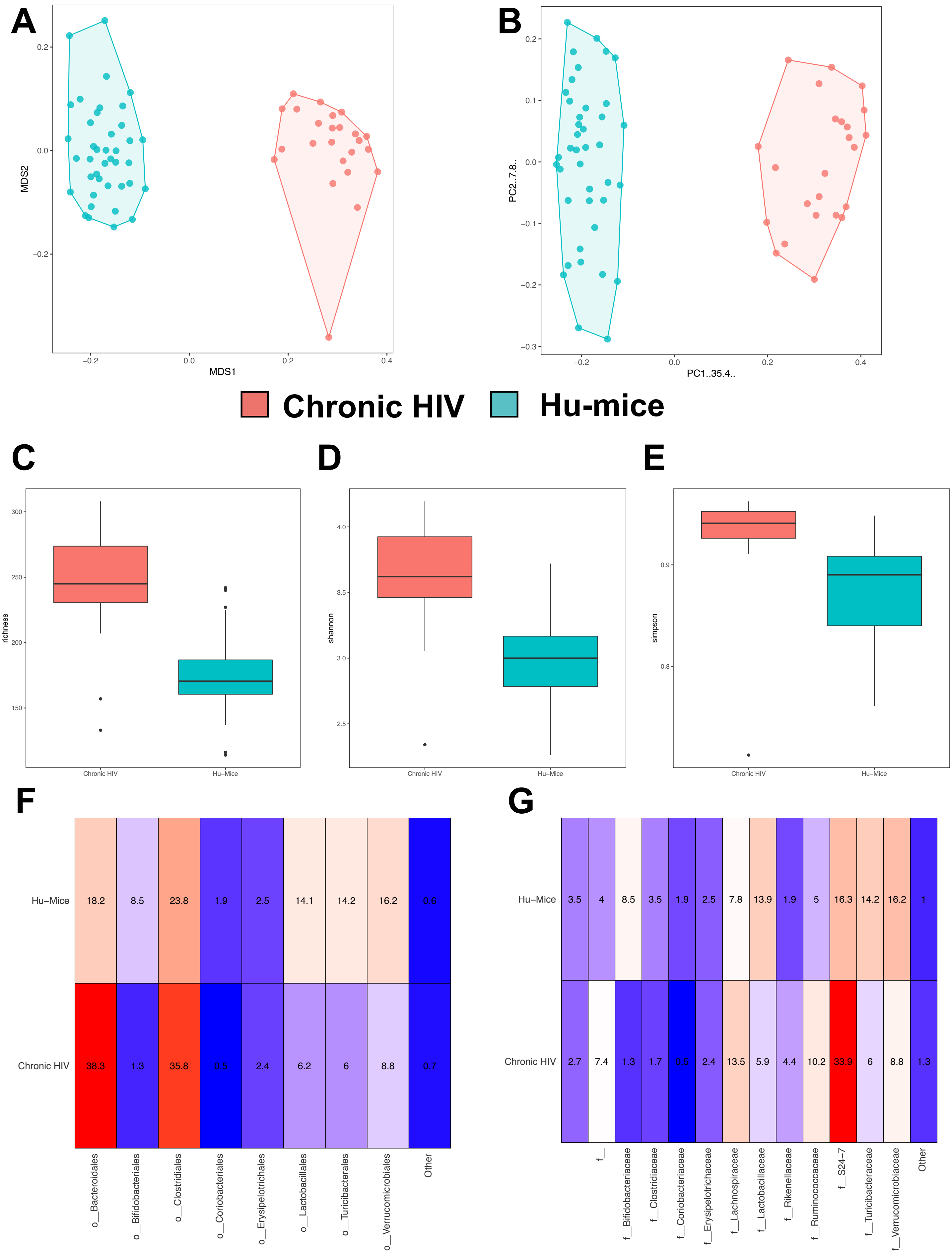

### Supplemental Figure HIV_Importance

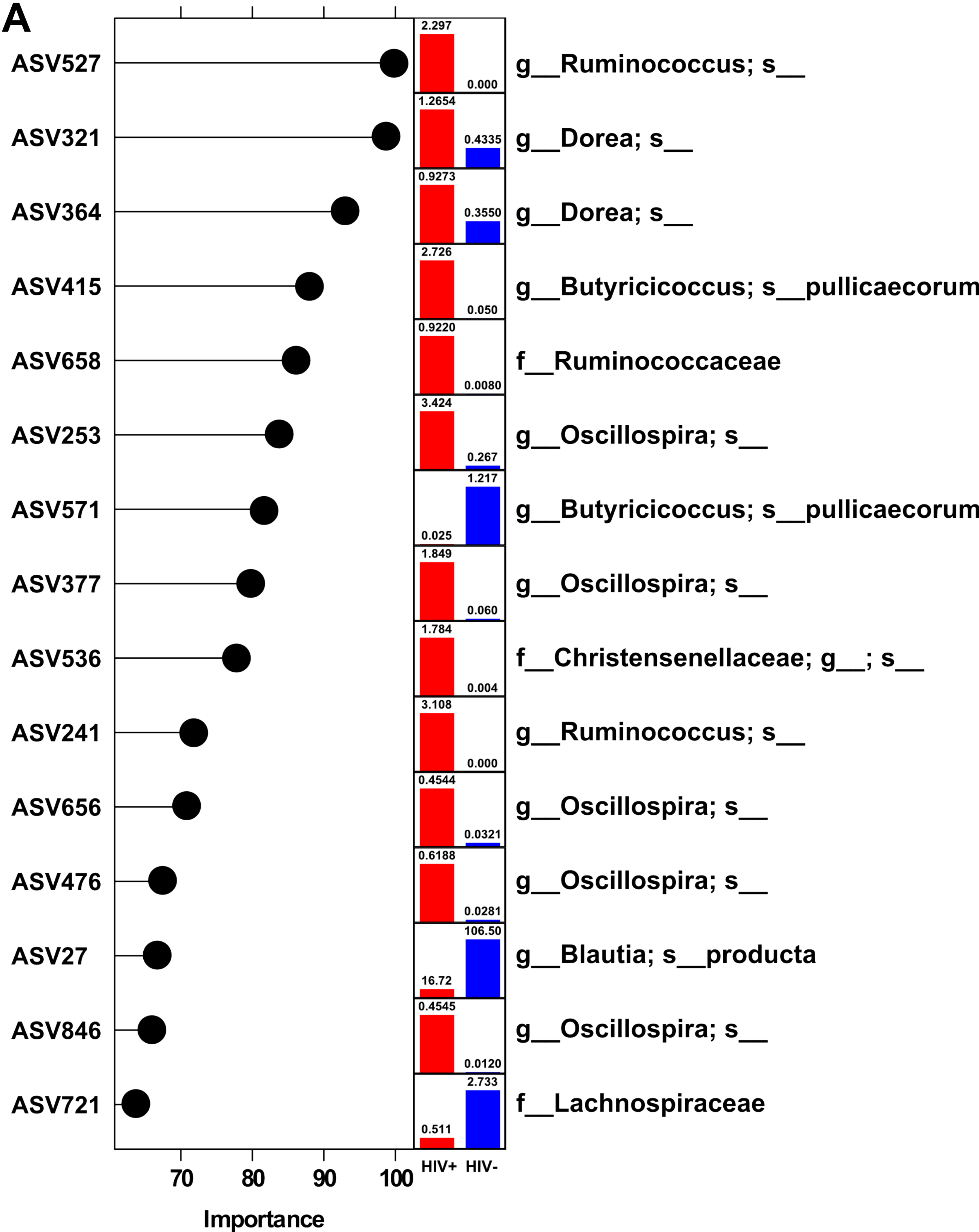

### Supplemental Figure HIV_Spleen

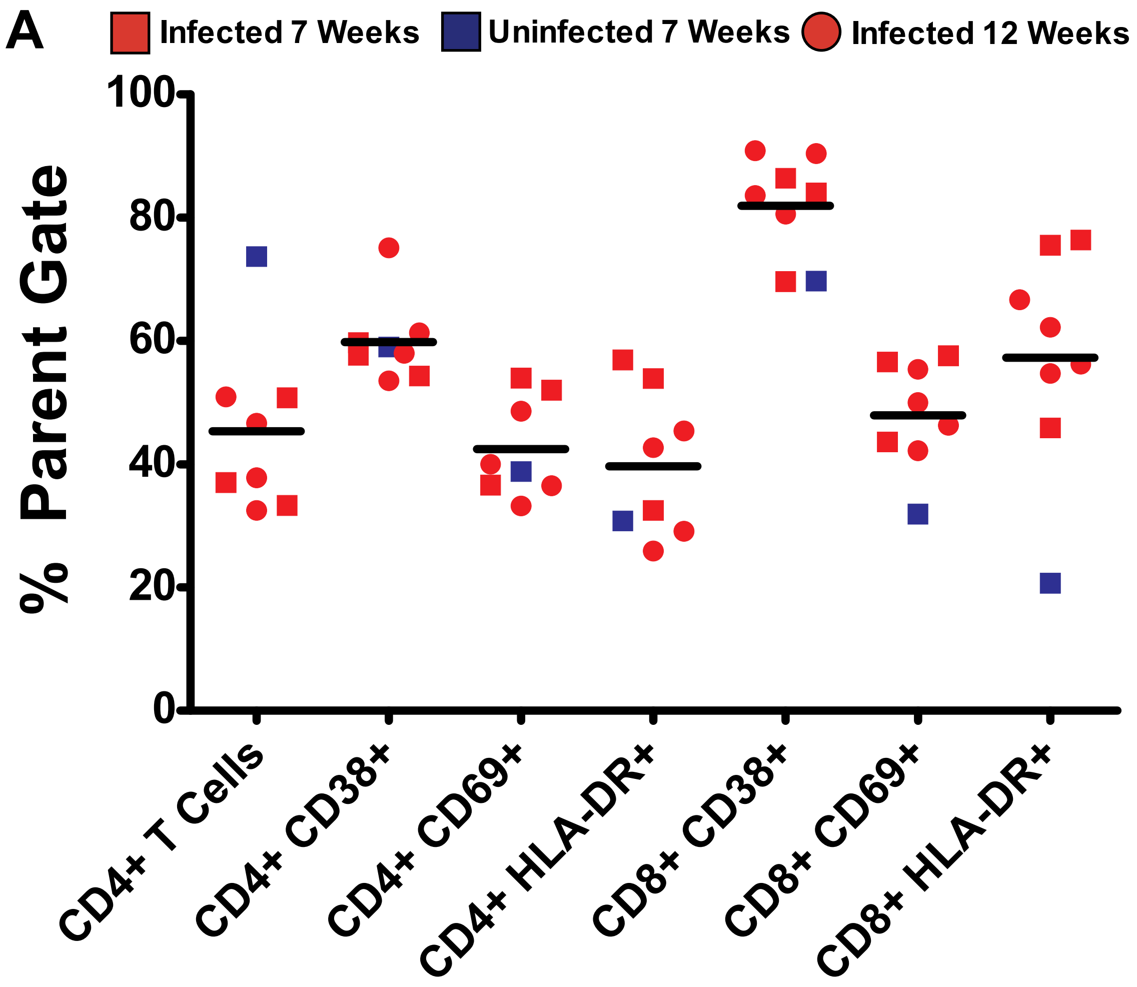

### Supplemental Figure Immune_Correlations

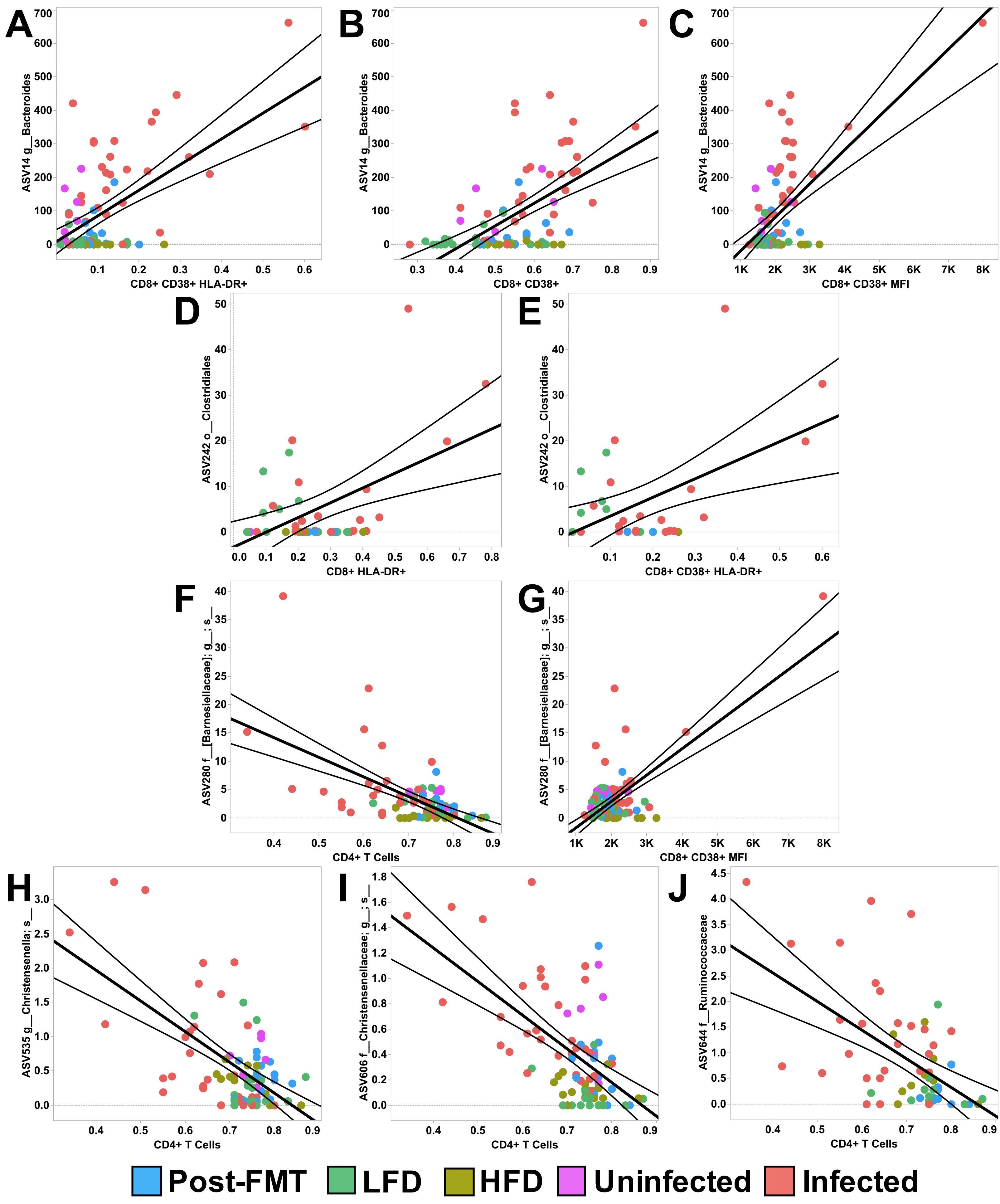

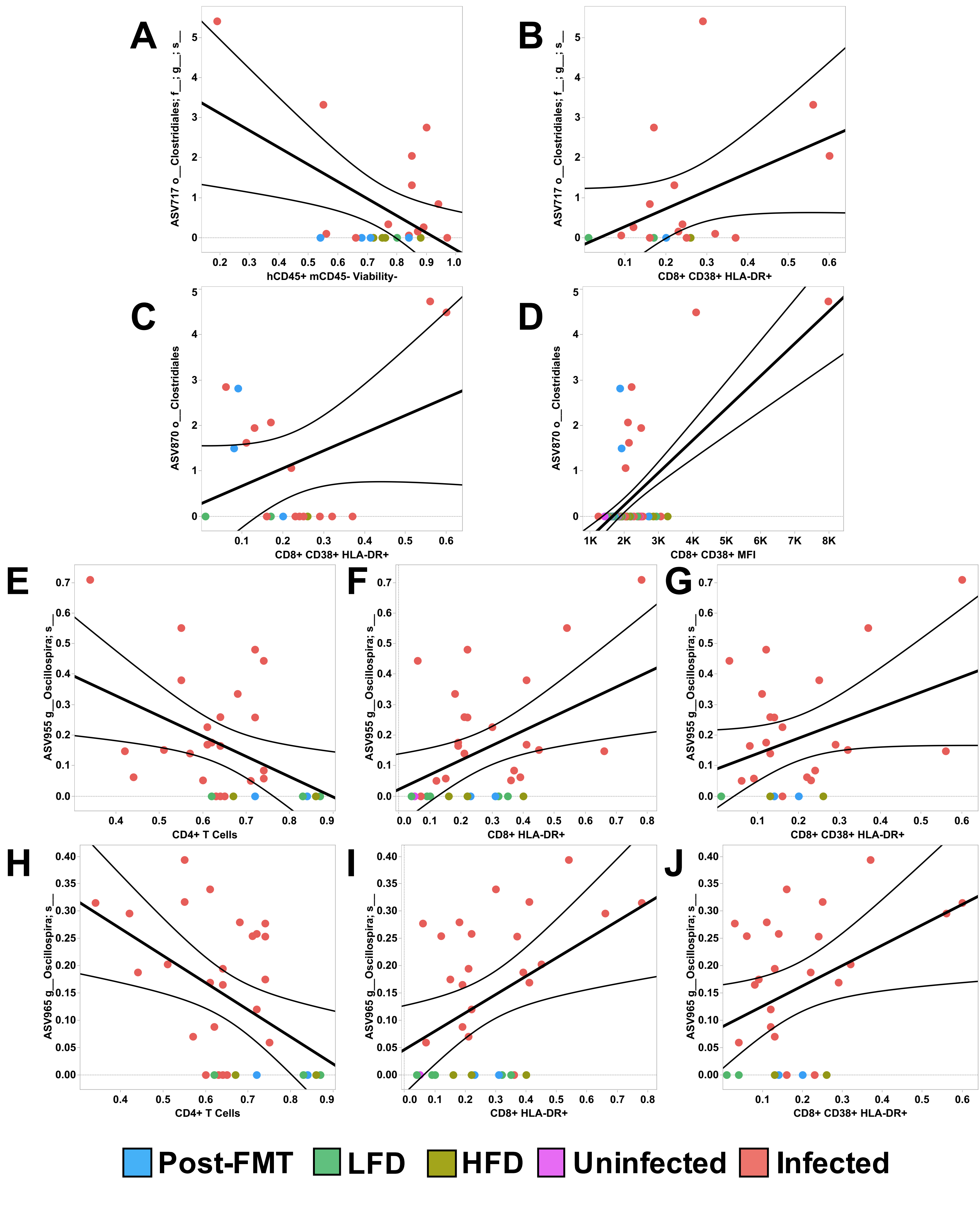
