## Supplemental Figure HIV_Composition for "Alterations in the gut microbiome by HIV-1 infection or a high-fat diet associates with systemic immune activation and inflammation in double humanized-BLT mice"

**A** Bifidobacteriaceae

| Infected | Post-FMT | 1 | 2 | 3 | 4 | 5 | 6 | 7 | 7S | 8 | 9 | 10 | 11 | 12S |
|---|---|---|---|---|---|---|---|---|---|---|---|---|---|---|
| 1292 | 1.5% | 32.2% | 20.7% | | 0.0% | 6.7% | 2.1% | 1.4% | | 7.0% | 0.4% | | | 0.0% |
| 1293 | 0.0% | 16.8% | 19.3% | | 28.1% | 3.6% | 11.3% | 0.5% | | 0.3% | 0.0% | | | 0.1% |
| 1294 | 0.0% | 1.1% | 6.1% | | 10.8% | 0.0% | 0.0% | 0.0% | 0.0% | | | | | |
| 1295 | 0.0% | 27.9% | 26.1% | | 5.5% | 5.8% | 5.9% | 0.7% | | 7.1% | 0.5% | | | 0.0% |
| 1311 | 0.8% | | | | | | | | | | | | | |
| 1316 | 0.1% | 0.0% | 0.0% | | 4.8% | 1.9% | 5.8% | 1.5% | 0.0% | | | | | |
| 1318 | 0.0% | 0.0% | 0.1% | | 4.6% | 1.9% | 3.5% | 9.4% | | 8.8% | 4.1% | | | 0.0% |
| 1321 | 0.0% | 21.7% | 24.5% | | 25.4% | 7.9% | 13.2% | 0.6% | 14.0% | | | | | |
| 1322 | 0.8% | 2.3% | 0.8% | | | | | | | | | | | |
| 1323 | 14.3% | 12.3% | 14.8% | | 5.4% | 1.2% | 3.7% | 0.0% | | | 0.0% | | | |
| 1324 | 3.5% | 3.4% | 8.8% | | | | | | | | | | | |
| 1325 | 16.2% | 11.5% | 16.3% | | 12.8% | 9.9% | 19.1% | 6.8% | | 1.3% | | | | |
| **Uninfected** | **Post-FMT** | **1** | **2** | **3** | **4** | **5** | **6** | **7** | **7S** | **8** | **9** | **10** | **11** | **12S** |
| 1317 | 0.0% | 0.0% | 0.0% | | 0.0% | 0.0% | 0.0% | 0.0% | 0.0% | | | | | |
| 1319 | 0.1% | 0.0% | 0.0% | | 0.0% | 0.0% | 0.0% | 0.0% | | 0.0% | 0.0% | | | |

**B** Rikenellaceae

| Infected | Post-FMT | 1 | 2 | 3 | 4 | 5 | 6 | 7 | 7S | 8 | 9 | 10 | 11 | 12S |
|---|---|---|---|---|---|---|---|---|---|---|---|---|---|---|
| 1292 | 0.6% | 0.8% | 1.2% | | 0.8% | 1.3% | 1.9% | 2.2% | | 3.3% | 2.3% | | | 1.5% |
| 1293 | 0.7% | 0.7% | 1.9% | | 1.2% | 4.0% | 2.7% | 4.4% | | 7.1% | 6.1% | | | 6.3% |
| 1294 | 1.3% | 1.1% | 0.7% | | 1.3% | 2.3% | 3.7% | 2.8% | 5.1% | | | | | |
| 1295 | 1.0% | 0.8% | 1.3% | | 2.4% | 1.5% | 2.7% | 2.7% | | 3.0% | 3.1% | | | 2.3% |
| 1311 | 0.2% | | | | | | | | | | | | | |
| 1316 | 1.1% | 2.8% | 2.0% | | 1.2% | 1.8% | 1.9% | 3.5% | 4.4% | | | | | |
| 1318 | 0.6% | 1.4% | 1.0% | | 1.4% | 1.2% | 1.5% | 1.8% | | 3.0% | 1.5% | | | 3.3% |
| 1321 | 0.2% | 0.8% | 0.5% | | 1.5% | 2.1% | 3.1% | 3.6% | 2.5% | | | | | |
| 1322 | 1.6% | 3.1% | 4.4% | | | | | | | | | | | |
| 1323 | 0.3% | 1.3% | 1.7% | | 1.6% | 2.7% | 2.7% | 4.9% | | | 4.7% | | | |
| 1324 | 1.4% | 2.4% | 1.1% | | | | | | | | | | | |
| 1325 | 1.0% | 1.8% | 2.2% | | 2.4% | 2.2% | 1.9% | 1.8% | | 6.1% | | | | |
| **Uninfected** | **Post-FMT** | **1** | **2** | **3** | **4** | **5** | **6** | **7** | **7S** | **8** | **9** | **10** | **11** | **12S** |
| 1317 | 0.7% | 1.4% | 0.6% | | 1.6% | 1.5% | 2.6% | 2.3% | 3.3% | | | | | |
| 1319 | 0.6% | 1.6% | 1.2% | | 1.8% | 2.6% | 2.6% | 4.0% | | 4.4% | 3.0% | | | |

**C** Staphylococcaceae

| Infected | Post-FMT | 1 | 2 | 3 | 4 | 5 | 6 | 7 | 7S | 8 | 9 | 10 | 11 | 12S |
|---|---|---|---|---|---|---|---|---|---|---|---|---|---|---|
| 1292 | 0.0% | 0.0% | 0.0% | | 0.6% | 0.0% | 0.0% | 0.0% | | 0.2% | 0.1% | | | 0.2% |
| 1293 | 0.0% | 0.0% | 0.0% | | 0.1% | 0.1% | 0.2% | 0.0% | | 0.2% | 0.1% | | | 0.4% |
| 1294 | 0.0% | 0.0% | 0.1% | | 0.0% | 0.1% | 0.5% | 0.0% | 0.0% | | | | | |
| 1295 | 0.0% | 0.0% | 0.0% | | 0.0% | 0.0% | 0.0% | 0.0% | | 0.0% | 0.1% | | | 0.3% |
| 1311 | 0.0% | | | | | | | | | | | | | |
| 1316 | 0.0% | 0.0% | 0.0% | | 0.0% | 0.0% | 0.0% | 0.0% | 0.0% | | | | | |
| 1318 | 0.0% | 0.1% | 0.0% | | 0.0% | 0.0% | 0.0% | 0.0% | | 0.1% | 0.1% | | | 0.2% |
| 1321 | 0.0% | 0.0% | 0.0% | | 0.0% | 0.0% | 0.0% | 0.0% | 0.0% | | | | | |
| 1322 | 0.4% | 0.0% | 0.0% | | | | | | | | | | | |
| 1323 | 0.1% | 0.0% | 0.0% | | 0.0% | 0.0% | 0.0% | 0.0% | | | 0.4% | | | |
| 1324 | 0.0% | 0.0% | 0.0% | | | | | | | | | | | |
| 1325 | 0.2% | 0.0% | 0.0% | | 0.0% | 0.0% | 0.0% | 0.0% | | 0.0% | | | | |
| **Uninfected** | **Post-FMT** | **1** | **2** | **3** | **4** | **5** | **6** | **7** | **7S** | **8** | **9** | **10** | **11** | **12S** |
| 1317 | 0.1% | 0.0% | 0.1% | | 0.1% | 0.0% | 0.0% | 0.0% | 0.1% | | | | | |
| 1319 | 0.0% | 0.0% | 0.0% | | 0.0% | 0.0% | 0.0% | 0.0% | | 0.1% | 0.0% | | | |

**D** Lactobacillaceae

| Infected | Post-FMT | 1 | 2 | 3 | 4 | 5 | 6 | 7 | 7S | 8 | 9 | 10 | 11 | 12S |
|---|---|---|---|---|---|---|---|---|---|---|---|---|---|---|
| 1292 | 0.2% | 3.8% | 0.4% | | 0.0% | 0.0% | 0.0% | 0.0% | | 0.0% | 0.0% | | | 0.0% |
| 1293 | 0.1% | 0.4% | 0.8% | | 0.0% | 0.2% | 0.1% | 0.0% | | 0.0% | 0.0% | | | 0.0% |
| 1294 | 0.0% | 0.0% | 0.0% | | 0.0% | 0.0% | 0.0% | 0.0% | 0.0% | | | | | |
| 1295 | 0.0% | 0.2% | 0.0% | | 0.0% | 0.0% | 0.0% | 0.0% | | 0.0% | 0.0% | | | 0.0% |
| 1311 | 0.0% | | | | | | | | | | | | | |
| 1316 | 32.9% | 24.3% | 14.1% | | 2.6% | 6.6% | 10.0% | 3.5% | 1.3% | | | | | |
| 1318 | 24.0% | 24.7% | 20.4% | | 13.4% | 29.3% | 20.4% | 26.9% | | 11.4% | 2.5% | | | 0.6% |
| 1321 | 0.0% | 0.0% | 0.3% | | 0.0% | 0.0% | 0.0% | 0.1% | 1.5% | | | | | |
| 1322 | 0.5% | 9.7% | 3.8% | | | | | | | | | | | |
| 1323 | 0.0% | 0.0% | 0.4% | | 0.0% | 0.0% | 0.1% | 0.0% | | | 16.3% | | | |
| 1324 | 0.0% | 0.0% | 0.4% | | | | | | | | | | | |
| 1325 | 0.0% | 0.2% | 0.0% | | 0.0% | 0.0% | 0.1% | 0.2% | | 0.0% | | | | |
| **Uninfected** | **Post-FMT** | **1** | **2** | **3** | **4** | **5** | **6** | **7** | **7S** | **8** | **9** | **10** | **11** | **12S** |
| 1317 | 18.7% | 34.8% | 38.0% | | 17.4% | 31.1% | 3.1% | 0.4% | 0.4% | | | | | |
| 1319 | 3.6% | 28.0% | 54.9% | | 20.3% | 22.1% | 18.9% | 8.0% | | 1.6% | 6.5% | | | |

**E** Turicibacteraceae

| Infected | Post-FMT | 1 | 2 | 3 | 4 | 5 | 6 | 7 | 7S | 8 | 9 | 10 | 11 | 12S |
|---|---|---|---|---|---|---|---|---|---|---|---|---|---|---|
| 1292 | 0.2% | 3.8% | 0.4% | | 0.0% | 0.0% | 0.0% | 0.0% | | 0.0% | 0.0% | | | 0.0% |
| 1293 | 0.1% | 0.4% | 0.8% | | 0.0% | 0.2% | 0.1% | 0.0% | | 0.0% | 0.0% | | | 0.0% |
| 1294 | 0.0% | 0.0% | 0.0% | | 0.0% | 0.0% | 0.0% | 0.0% | 0.0% | | | | | |
| 1295 | 0.0% | 0.2% | 0.0% | | 0.0% | 0.0% | 0.0% | 0.0% | | 0.0% | 0.0% | | | 0.0% |
| 1311 | 0.0% | | | | | | | | | | | | | |
| 1316 | 32.9% | 24.3% | 14.1% | | 2.6% | 6.6% | 10.0% | 3.5% | 1.3% | | | | | |
| 1318 | 24.0% | 24.7% | 20.4% | | 13.4% | 29.3% | 20.4% | 26.9% | | 11.4% | 2.5% | | | 0.6% |
| 1321 | 0.0% | 0.0% | 0.3% | | 0.0% | 0.0% | 0.0% | 0.1% | 1.5% | | | | | |
| 1322 | 0.5% | 9.7% | 3.8% | | | | | | | | | | | |
| 1323 | 0.0% | 0.0% | 0.4% | | 0.0% | 0.0% | 0.1% | 0.0% | | | 16.3% | | | |
| 1324 | 0.0% | 0.0% | 0.4% | | | | | | | | | | | |
| 1325 | 0.0% | 0.2% | 0.0% | | 0.0% | 0.0% | 0.1% | 0.2% | | 0.0% | | | | |
| **Uninfected** | **Post-FMT** | **1** | **2** | **3** | **4** | **5** | **6** | **7** | **7S** | **8** | **9** | **10** | **11** | **12S** |
| 1317 | 18.7% | 34.8% | 38.0% | | 17.4% | 31.1% | 3.1% | 0.4% | 0.4% | | | | | |
| 1319 | 3.6% | 28.0% | 54.9% | | 20.3% | 22.1% | 18.9% | 8.0% | | 1.6% | 6.5% | | | |

**F** Lachnospiraceae

| Infected | Post-FMT | 1 | 2 | 3 | 4 | 5 | 6 | 7 | 7S | 8 | 9 | 10 | 11 | 12S |
|---|---|---|---|---|---|---|---|---|---|---|---|---|---|---|
| 1292 | 17.1% | 6.3% | 8.6% | | 13.5% | 16.2% | 8.6% | 10.6% | | 12.7% | 17.7% | | | 16.2% |
| 1293 | 12.9% | 11.1% | 7.2% | | 6.2% | 12.8% | 9.4% | 12.8% | | 10.2% | 11.6% | | | 9.5% |
| 1294 | 15.3% | 19.6% | 11.8% | | 10.0% | 12.2% | 11.1% | 25.3% | 13.8% | | | | | |
| 1295 | 19.1% | 10.6% | 7.6% | | 11.2% | 11.3% | 9.7% | 11.3% | | 12.9% | 10.6% | | | 8.1% |
| 1311 | 30.8% | | | | | | | | | | | | | |
| 1316 | 4.5% | 9.5% | 8.8% | | 11.6% | 10.4% | 7.6% | 14.3% | 11.8% | | | | | |
| 1318 | 6.9% | 10.8% | 9.2% | | 8.5% | 7.7% | 11.7% | 6.8% | | 7.4% | 14.0% | | | 13.4% |
| 1321 | 17.5% | 10.2% | 10.6% | | 5.3% | 6.5% | 8.4% | 14.2% | 11.3% | | | | | |
| 1322 | 20.8% | 13.1% | 10.5% | | | | | | | | | | | |
| 1323 | 20.3% | 15.8% | 9.2% | | 13.6% | 20.1% | 10.2% | 11.3% | | | 14.6% | | | |
| 1324 | 17.7% | 12.9% | 13.5% | | | | | | | | | | | |
| 1325 | 17.2% | 13.9% | 8.9% | | 10.9% | 12.1% | 7.2% | 12.1% | | 10.3% | | | | |
| **Uninfected** | **Post-FMT** | **1** | **2** | **3** | **4** | **5** | **6** | **7** | **7S** | **8** | **9** | **10** | **11** | **12S** |
| 1317 | 8.2% | 5.6% | 6.2% | | 8.0% | 6.4% | 9.1% | 12.4% | 10.4% | | | | | |
| 1319 | 19.6% | 8.7% | 5.4% | | 9.1% | 7.2% | 9.1% | 10.7% | | 14.1% | 15.6% | | | |

**G** Ruminococcaceae

| Infected | Post-FMT | 1 | 2 | 3 | 4 | 5 | 6 | 7 | 7S | 8 | 9 | 10 | 11 | 12S |
|---|---|---|---|---|---|---|---|---|---|---|---|---|---|---|
| 1292 | 9.6% | 8.4% | 12.3% | | 19.7% | 18.4% | 15.2% | 20.5% | | 24.2% | 19.1% | | | 11.6% |
| 1293 | 2.6% | 7.8% | 13.8% | | 10.1% | 16.5% | 14.2% | 14.2% | | 15.0% | 13.5% | | | 20.3% |
| 1294 | 5.4% | 9.9% | 10.6% | | 16.2% | 13.8% | 12.1% | 16.1% | 17.8% | | | | | |
| 1295 | 7.9% | 16.2% | 17.0% | | 16.4% | 18.0% | 20.1% | 20.4% | | 19.7% | 18.2% | | | 4.8% |
| 1311 | 3.0% | | | | | | | | | | | | | |
| 1316 | 2.2% | 11.6% | 6.6% | | 12.4% | 14.6% | 16.1% | 13.7% | 18.3% | | | | | |
| 1318 | 6.9% | 11.2% | 9.2% | | 10.2% | 10.7% | 13.4% | 8.7% | | 17.3% | 17.5% | | | 17.5% |
| 1321 | 6.2% | 10.8% | 13.9% | | 12.5% | 14.7% | 10.9% | 13.1% | 9.5% | | | | | |
| 1322 | 6.7% | 12.7% | 17.0% | | | | | | | | | | | |
| 1323 | 7.1% | 16.4% | 15.4% | | 16.1% | 18.6% | 14.4% | 21.3% | | | 8.9% | | | |
| 1324 | 7.2% | 13.5% | 17.0% | | | | | | | | | | | |
| 1325 | 4.1% | 16.9% | 17.2% | | 16.5% | 16.0% | 13.8% | 19.2% | | 17.5% | | | | |
| **Uninfected** | **Post-FMT** | **1** | **2** | **3** | **4** | **5** | **6** | **7** | **7S** | **8** | **9** | **10** | **11** | **12S** |
| 1317 | 8.0% | 6.0% | 7.8% | | 9.6% | 7.1% | 10.8% | 13.0% | 11.7% | | | | | |
| 1319 | 8.5% | 10.9% | 7.5% | | 8.9% | 8.5% | 9.4% | 10.1% | | 15.0% | 17.8% | | | |

**H** Desulfovibrionaceae

| Infected | Post-FMT | 1 | 2 | 3 | 4 | 5 | 6 | 7 | 7S | 8 | 9 | 10 | 11 | 12S |
|---|---|---|---|---|---|---|---|---|---|---|---|---|---|---|
| 1292 | 0.2% | 0.4% | 0.2% | | 1.7% | 0.8% | 1.5% | 1.0% | | 0.5% | 0.7% | | | 1.1% |
| 1293 | 0.9% | 0.4% | 0.8% | | 0.5% | 0.8% | 0.9% | 1.3% | | 0.7% | 0.9% | | | 1.2% |
| 1294 | 1.9% | 0.8% | 0.8% | | 0.3% | 1.9% | 1.8% | 0.5% | 0.9% | | | | | |
| 1295 | 0.4% | 0.3% | 0.6% | | 0.5% | 0.4% | 0.8% | 0.8% | | 0.6% | 0.7% | | | 0.3% |
| 1311 | 0.9% | | | | | | | | | | | | | |
| 1316 | 0.5% | 0.3% | 0.9% | | 0.7% | 1.0% | 0.5% | 0.9% | 1.1% | | | | | |
| 1318 | 0.7% | 0.9% | 0.7% | | 0.2% | 0.4% | 0.4% | 0.4% | | 0.5% | 0.5% | | | 1.7% |
| 1321 | 0.7% | 0.4% | 0.2% | | 0.4% | 0.7% | 0.7% | 2.1% | 1.1% | | | | | |
| 1322 | 0.7% | 1.0% | 2.3% | | | | | | | | | | | |
| 1323 | 0.5% | 0.7% | 0.7% | | 0.8% | 0.7% | 0.9% | 1.3% | | | 0.6% | | | |
| 1324 | 0.8% | 1.8% | 2.6% | | | | | | | | | | | |
| 1325 | 0.4% | 0.8% | 0.4% | | 0.4% | 0.5% | 0.4% | 0.4% | | 1.0% | | | | |
| **Uninfected** | **Post-FMT** | **1** | **2** | **3** | **4** | **5** | **6** | **7** | **7S** | **8** | **9** | **10** | **11** | **12S** |
| 1317 | 0.5% | 0.9% | 0.2% | | 0.2% | 0.4% | 0.8% | 1.0% | 1.1% | | | | | |
| 1319 | 0.4% | 0.6% | 0.5% | | 0.5% | 0.6% | 0.6% | 0.6% | | 0.5% | 0.3% | | | |
